# A robust and integrated framework for cross-platform adaptation of epigenetic clocks in cell-free DNA sequencing

**DOI:** 10.1101/2025.11.27.690895

**Authors:** Guangyu Li, Weilai Huang, Xuechen Zhao, Jun Wu, Yuanfang Guo, Liying Chen, Xingqi Cao, Zipeng Yang, Siqi Jiang, Bin Hu, Yijun Wang, Dinson Tan, Vicky Tong, Changdong Tang, Xiaowen Feng, Xiaomin Hu, Chuan Ouyang, Guangyu Zhou

## Abstract

**Background.:** Circulating cell-free DNA (cfDNA or ccfDNA) methylation sequencing holds promise for developing epigenetic aging clocks in minimally invasive aging assessment applications. However, current clocks—primarily trained on array-based data—do not readily generalize to methylation profiles of cfDNA, due to stochasticity and uncertainty arising from the limited input amounts of cfDNA and technical characteristics of high-throughput sequencing (HTS) platform. Due to lack of training to correct this uncertainty, direct application of legacy clocks to HTS data will inevitably produce unreliable estimates, introduce predictive discordance and undermine the trustworthiness of epigenetic biomarkers. Despite this urgent challenge, a systematic, model-agnostic framework for adapting existing epigenetic clocks to HTS-based cfDNA data remains lacking.

**Methods.:** Here, we generated a dedicated benchmark dataset comprising paired cfDNA and genomic DNA (gDNA), which was replicated and profiled across two methylation arrays and two targeted HTS platforms. We evaluated 53 epigenetic clocks for CpG coverage, reproducibility, cross-platform consistency, and age prediction accuracy. Further, we systematically explored and tested multiple adaptation strategies including depth filtering, beta-value imputation, and transfer learning via model distillation aiming at improving performance and clinical concordance of legacy epigenetic clocks on cfDNA HTS datasets.

**Results.:** Our results show that inherent technical noise of HTS platforms compromises diagnostic precision, presenting as a significant burden for applying legacy clocks on cfDNA HTS data. However, this technical limitation can be systematically neutralized. By enforcing stringent sequencing depth protocols (≥10× ideally 20×), employing robust algorithmic stabilization (L2-heavy clocks and imputation) and transfer learning, we can effectively isolate genuine physiological aging signals from technical artifacts. Ultimately, an adaptation pipeline incorporating all these methods effectively improved age prediction accuracy and, closed the gap between epigenetically induced aging status and clinically assessed realities. Besides, the pipeline demonstrated improved diagnostic sensitivity for neurodegenerative conditions such as amyotrophic lateral sclerosis, establishing a reliable foundation for non-invasive clinical monitoring.

**Conclusions.:** This study provided a comprehensive framework and practical guidelines for adapting epigenetic clocks to HTS-based cfDNA data, paved the path for more general application of cfDNA-based aging assessment and liquid biopsies.

## Introduction

Recent studies increasingly advocate using biological age (BA), rather than chronological age (CA), as a more informative indicator of an individual’s physiological aging state. BA is estimated from biomarkers of aging (BoAs) through mathematical models known as aging clocks [1,2]. Among diverse BoAs, epigenetic clocks—based on the methylome (DNA methylation profiles)—have emerged as leading tools due to their outstanding predictive accuracy [3–5], achievable even with simple models [4,6,7]. Consequently, epigenetic clocks are now widely used in health monitoring, age-related disease assessment, and longevity intervention studies [2,3,8].

Circulating cell-free DNA (cfDNA or ccfDNA), primarily found in body fluids, enables minimally invasive testing for early cancer detection and disease monitoring [9]. Its utility in epigenetic aging clocks is being actively explored [10–12]. Unlike genomic DNA (gDNA), methylation profiling of cfDNA often required high-throughput sequencing (HTS) technologies over array technologies (e.g. Illumina MethylationEPICv2 BeadChip) due to several biological and technical limitations, including high tissue-of-origin heterogeneity, short fragment sizes, low input (0.1–10 ng, compared with 200 ng recommended for arrays), high dynamic range, requirement to resolve intra-sample methylation heterogeneity [9,13–15]. Though some limitations (e.g. low input) can be technically overcome, drawbacks (e.g. distorted beta-values) still hinder the clinical translation[16]. In practice, targeted sequencing based on hybridization probe capture is the primary choice for cfDNA methylation profiling [17–19].

This technological platform mismatch hampers the application of legacy epigenetic clocks, which are often trained on array-based gDNA data, to cfDNA data. The incompatibility is fundamentally rooted in divergent data structures: microarrays generate continuous beta-values from fluorescent signal intensities with predictable variance, whereas HTS yields discrete, count-based methylation ratios that are inherently heteroscedastic. This data-level discordance threatens using legacy clocks in cross-platform research, as directly exposing legacy clocks to the unseen technical and biological variance inevitably triggers unpredictable model behavior [15,20,21]. In cross-sectional studies, this may widen the discordance among different clocks, resulting in loss of accuracy; in longitudinal studies, this may contribute to false identification of aging or disease trajectories, and most critically, reducing the trustworthiness of related epigenetic biomarkers. Hence, closing this platform gap is critical in cfDNA-based epigenetic aging assessments.

To bridge this gap, two methodological paradigms have emerged. First, *de novo* HTS models, such as BS-clock and timeseqage [15,22], are trained exclusively on sequencing datasets to bypass platform-specific biases. However, they typically identify novel HTS-specific feature sets rather than retaining the established biological relevance of legacy array-trained clocks. Second, adaptation strategies aim to harmonize array-trained models with HTS data, mainly categorized into data-centric approaches (e.g. batch correction [23–25] and domain adaptation [26,27]) and model-transfer frameworks (e.g. Transfer Elastic Net [28]). However, the former ones may introduce technical artifacts [29], while the latter ones are fundamentally model-specific, requiring architectural transparency, even original training data that have no guaranteed accessibility. Some recent robust feature-selection methods attempt to identify consensus features across platforms [30], but their reliance on cell-type-specific tuning is largely incompatible with the intrinsic cell type heterogeneity of cfDNA. Furthermore, the scarcity of systematic benchmark resources featuring paired array/HTS technical replicates has hindered the establishment of standardized experimental and bioinformatic protocols — a gap particularly acute for cfDNA, where platform-specific biases readily confound biological aging signals. Therefore, a systematic, model-agnostic framework capable for benchmarking and adapting legacy array-trained clocks for high-precision, HTS-based cfDNA analysis, is in need.

To systematically address the platform gap, we established a controlled benchmarking framework using a matched dataset of cfDNA and gDNA profiled by both HTS and array-based methylation technologies. This design enabled quantitative assessment of clock accuracy, precision, and reproducibility across DNA types and platforms, and allowed us to decompose and correct sources of variance, including coverage depth, CpG missingness, and platform biases. Through structured evaluation of these experimental and computational factors, we identified recurrent bias patterns and failure modes arising from cross-platform transfer of epigenetic clocks. Based on these insights, we developed and validated an end-to-end adaptation pipeline integrating experimental design guidance, data harmonization, feature recalibration, and machine-learning-based correction. Collectively, this work provides a systematic benchmarking resource and a validated methodological framework for robust application of epigenetic clocks to HTS-derived cfDNA data. Establishing such a high-fidelity digital representation of epigenetic information is critical for the clinical utility of liquid biopsies and the broader advancement of precision medicine, providing the necessary foundation for longitudinal health tracking and the data-driven optimization of individualized therapeutic interventions [31].

## Methods

### The SRRSH cohorts and ethics declaration

The ethics committee of the Sir Run Shaw Hospital (SRRSH, affiliated with Zhejiang University School of Medicine) approved this study (approval number 2024-0547). Written informed consent was obtained from all participants.

The recruitment and data collection for all volunteers in the cohort were conducted under strict protocols. The following criteria were applied for recruiting eligible volunteers: (1) healthy adults aged 20 - 84 years; (2) the subject has completed at least two routine health evaluations within the past 5 years that included complete blood count, urinalysis, fasting plasma glucose, lipid profile, liver-function tests, renal-function tests, and at least one set of imaging examinations: abdominal ultrasound (liver, gall-bladder, pancreas, spleen, both kidneys), chest radiograph, thyroid ultrasound, prostate ultrasound (men), uterus and adnexa ultrasound (women). All reported results must be within normal limits and free of any diagnosed disease; (3) no smoking history within 5 years before screening and agreement to remain abstinent from tobacco for 1 year after consent; (4) body-mass index 18.5-28.0 kg/m^2^; (5) without acute infection or active inflammation within 14 days before screening; (6) without cancer-related signs or symptoms (e.g., hemoptysis, rectal bleeding, altered bowel habit, hematuria, dysphagia, palpable mass) within 30 days before screening; (7) without previous organ transplantation or any non-autologous bone-marrow or stem-cell transplantation; (8) without blood transfusion within 7 days before screening; (9) without use of therapeutic drugs for any illness (nutritional supplements excluded) within 30 days before screening; (10) without any surgical procedure within 5 years before screening; (11) pregnant or breastfeeding women were excluded.

### Methylation array and capture panel selection

Four methylation profiling methods were included in this study, including two array-based methods and two HTS-based methods with different hybridization probe capture panels: Illumina Human Methylation Screening Array (referred to as MSA), Illumina Methylation EPIC v2.0 BeadArray (referred to as EPICv2), targeted sequencing with iGeneTech BisCap CpG Galaxy Panel (referred to as Galaxy) and targeted sequencing with Twist Human Methylome Panel (referred to as Twist). EPICv2 was included because it encompasses the largest CpG set among currently available methylation arrays, which serve as the basis for training most existing epigenetic clocks. MSA is a newer methylation array with a smaller CpG set than EPICv2, yet is claimed by the manufacturer to be suitable for epigenetic clock applications. Galaxy was included because it is designed as an alternative to EPICv2, which covers over 95% of its CpG sites. Lastly, Twist was included as a mature solution for HTS-based methylation profiling with appropriate CpG coverage in the current marketplace. Overall, these methods were selected to cover mature, widely used, and emerging methods in both methylation array and HTS categories.

### Experimental design

In total, 165 participants (100 % East Asian; aged 20 - 77 years old; 41.8 % female) were recruited as the SRRSH cohort. From this cohort, 24 subjects were randomly selected (22 - 77 years old; 54.2 % female) for investigating and evaluating the adaptation of epigenetic clocks for cfDNA-based methylation data, referred to as the SRRSH-24 cohort. Though no sample size calculations were conducted *a priori*, a previous study suggested a similar sample size was sufficient to identify 80% of reliable probes [32]. Their cfDNA was extracted for targeted DNA methylation sequencing, and buffy coat gDNA was extracted for array-based and HTS-based methylation profiling. This resulted in six datasets: gDNA_MSA, gDNA_EPICv2, gDNA_Galaxy, gDNA_Twist, cfDNA_Galaxy, and cfDNA_Twist. Two technical replicates per sample were generated via independent library preparations from the same DNA aliquot. Finally, the SRRSH-141 dataset contained Twist-profiled plasma cfDNA methylation data for the remaining 141 participants. The sample details are listed in Table S2.

### Epigenetic clock selection

A total of 57 epigenetic clocks were included in this study, including 53 array-trained and 4 HTS-trained clocks (Table S1). The 53 array-trained clocks include 47 integrated in the pyaging package (version 0.1.27) and 6 GP-age clocks (https://github.com/mirivar/GP-age, version 1.0). The 4 HTS-trained clocks include emseqage [10], timeseqage (https://github.com/patricktgriffin/TIME-Seq, Commit 140622f), BS-clock (https://github.com/hucongcong97/BS-clock, Commit 097b800) and komakiage [33]. Two clocks, BS-clock and komakiage, were not included in analyses beyond feature coverage assessment due to their overall low feature coverage in all six SRRSH-24 datasets.

### Biological sample pretreatment and storage

An 8-10 mL whole-blood sample was collected from each volunteer via venipuncture, after an overnight fast. One anticoagulant tube (Streck, Cell-Free DNA BCT 10ml) was immediately transported to the laboratory for subsequent treatment. The blood was centrifuged at 4°C (1900× g, 10 min) to obtain plasma and buffy coat. All biological samples were stored at −80°C for long-term preservation.

### DNA extraction

Cell-free DNA was extracted from 1.0 - 1.5 mL plasma using the QIAamp MinElute ccfDNA Mini Kit (Qiagen, Cat# 55204) following the manufacturer’s instructions. Genomic DNA was isolated from approximately 500 μL of thawed buffy coat using TIANamp Genomic DNA Kit (TIANGEN, Cat# DP304) following the manufacturer’s protocol. The concentration of purified DNA was quantified using a Thermo Fisher Qubit 4.0 fluorometer (Qubit™ dsDNA HS Assay Kit, Cat# Q32854).

### Library preparation and sequencing

For genomic DNA, 0.5% (by mass) of unmethylated lambda DNA and 0.05% (by mass) of CpG-methylated pUC19 DNA were spiked into 400-800 nanograms of buffy coat genomic DNA, followed by dilution to 130 μL of 10 mM Tris 0.1 mM EDTA, pH 8.0, following the manufacturer’s recommendations. Internal control DNA underwent the same fragmentation procedure as gDNA samples. The DNA mixture was then transferred to a Covaris microTUBE and sheared to an average size of 100-200 bp (peak = 150bp) using a Covaris™ S220 focused ultrasonicator (Covaris, MA, USA).

For cfDNA, unmethylated lambda DNA and CpG methylated pUC19 DNA were respectively sheared to an average size of 100-200 bp (peak = 150bp) in advance. Next, 0.5% (by mass) of fragmented unmethylated lambda DNA and 0.05% (by mass) of fragmented CpG methylated pUC19 DNA were added into 1-10 ng cfDNA samples.

Both cfDNA and sonicated gDNA samples, along with spike-in internal control DNA, were used to construct EM-Seq libraries with the NEBNext Enzymatic Methyl-seq Kit (New England Biolabs, Cat# E7120) following the manufacturer’s instructions. Libraries were quantified using a Thermo Fisher Qubit 4.0 fluorometer. Then capture-based human methylome enrichment was performed according to Twist’s (South San Francisco, USA) protocols for their Twist Human Methylome Panel (Cat# 105521), or according to iGeneTech’s (Beijing, China) protocols for their BisCap CpG Galaxy Panel (Cat# PE4000175). The final libraries were analyzed and quantified using TapeStation (Agilent Technologies). The whole-genome libraries were 150 bp paired-end sequenced using the Illumina NovaSeq X Plus sequencer on a 10B or 25B flow cell (Illumina) with 12.5 % PhiX spiked-in, to an average yield of 36.3Gbp raw bases per library (Fig S9).

### EPICv2 and MSA

Buffy coat genomic DNA samples were sent to Novogene (Tianjin, China) to be analyzed using the Illumina Infinium MethylationEPIC v2.0 BeadChip array (EPICv2; 937,691 probes), or sent to WeGene (Shenzhen, China) for profiling on the Infinium Methylation Screening array (MSA; 284,318 probes). For each technical replicate sample, 500 nanograms (for EPICv2) or 200 nanograms (for MSA) of genomic DNA were bisulfite treated using the EZ DNA Methylation Kit (Zymo Research, Cat# D5001) following the manufacturer’s recommendations for Infinium assays.

As the EPICv2 and MSA arrays can accommodate 8 and 48 samples per array, respectively, we included two replicates (2 replicates × 24 gDNA samples = 48 slots).

### Methylation array data processing

Raw IDAT files (from both EPICv2 and MSA) were processed using the `minfi` v1.54.1 according to established protocols. For rigorous quality control, following probes are removed: (1) located on sex chromosomes; or (2) detection p-value ≥ 0.01; or (3) probes known to harbor single nucleotide polymorphisms (SNPs) at or near the CpG site; or (4) with ambiguous mapping. Intra-array normalization of raw intensities was performed using the noob method.

### HTS methylation data processing

Raw paired-end FASTQ files were first downsampled to standardize sequencing depth at approximately 30 Gbp (1 Gbp = 10^9^ base pairs) to ensure fair comparisons. Subsequent steps include: (1) adapter removal (`fastp` v0.23.4), alignment (`bwa-meth` v0.2.7), sorting and indexing (`samtools` v1.21), duplicate removal (`samblaster` v 0.1.26), and beta-value extraction (`MethylDackel` v0.6.1). The `bwa-meth` alignment uses a combined reference comprising GRCh38, pUC19, and phage lambda sequences.

### Genomic coordinate harmonization and probe annotation

All in-house and public datasets were harmonized to hg38 genome coordinates using UCSC `LiftOver` whenever applicable. Probes defined in any of the array probe manifestation files (downloaded from manufacturers’ sites) were annotated based on their genome coordinates.

### Statistical analysis

Permutation tests were implemented using custom script with p-values calculated as the proportion of random shuffles yielding statistics more extreme than the observed data. This non-parametric approach requires no distributional assumptions and remains robust across various sample sizes. In multiple comparison scenarios, the p-values were adjusted using the Benjamini-Hochberg (BH) method.

### Public datasets

See Supplementary Information Appendix for details.

### Elastic net model training for independent verification of the correlation between number of CpGs and RD

A series of elastic net models with varying alpha and l1_ratio hyperparameters were trained on the dataset GSE55763 (N = 2711) using `ElasticNet` from the `scikit-learn` python package (version 1.6.1). The input features were limited to the 90,394 CpGs shared between GSE55763 and the SRRSH-24 datasets. Missing values, if any, were imputed using KNN.

### Simulating impact of sequencing depth based on binomial stochasticity

To simulate the stochasticity of observed value beta’ distributed near a latent truth beta, we used the following formula:

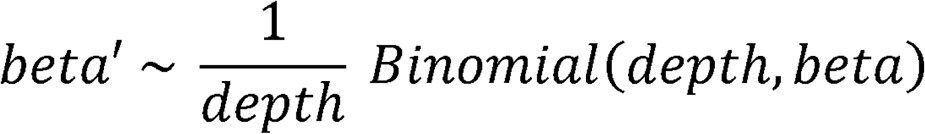

Where:

depth is the sequencing for under investigation.

beta is the beta-value observed in the dataset.

The final statistical properties of MAE and RD accounting for this stochasticity were evaluated after 1,000 repeats to reduce randomness.

### Simulating impact of sequencing depth based on beta resolution

For each sequencing depth, the beta values are rounded by the below formula, to simulate the precision loss of beta values under a finite sequencing depth:

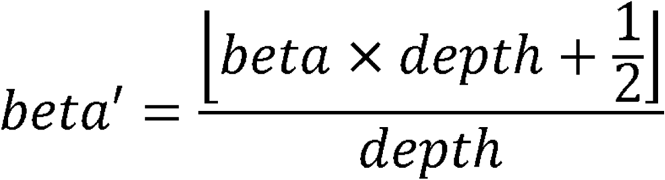

depth is the sequencing depth for investigation.

beta is the beta-value observed in the dataset.

⌊·⌋ is the floor function.

### Minimal sequencing investigation based on downsampling

5 samples with initial sequencing result size ≥50Gb were pre-selected, then downsampled to 0.025x to 0.9x (namely 0.025, 0.05, 0.1, 0.2, 0.3, 0.4, 0.5, …, 0.9) before biological age estimation.

### Unreliable beta-value imputation at different cut-off depths

This strategy encompasses three consecutive steps: (1) Filling original missing values; (2) Beta values of value 0 or 1 with corresponding sequencing depth under chosen threshold was set to missing values (NaNs). (3) Imputation of missing values set by step (2).

### Transfer learning of classic epigenetic models

Each HTS dataset was first PCA-transformed using PCA models independently trained from public dataset GSE55763 (N = 2711) with an output dimensionality of 2500. The student elastic net model was trained using PCA-transformed dataset using the corresponding teacher model predictions as the supervision target (soft labels) and a 5-fold cross-validation. The hyperparameters (`alpha` and `l1_ratio`) were optimized during training using an independent 5-fold cross-validation.

### The DF-IM-TL Adaptation Pipeline

To overcome the “platform gap” between continuous array signals and discrete HTS counts, we developed the DF-IM-TL pipeline: DF: CpG sites with sequencing coverage below 10× were excluded to mitigate technical stochasticity. IM: Extreme beta-values (approaching 0 or 1) arising from low-coverage sites were imputed. TL: A teacher-student distillation model was employed to align HTS-derived predictions with array-based benchmarks.

### Adaptation baseline methods comparison

Three adaptation methods were evaluated as baselines for comparison with the DF-IM-TL pipeline. Quantile mapping was implemented using custom script, which replaces values in the adapting dataset with values at respective quantile in the reference dataset. ComBat was performed using neuroCombat (version 0.2.12) package. Correlation alignment (CORAL) was implemented using custom script, was performed after a truncated singular value decomposition for dimensionality reduction.

### MAPLE training

Besides the pretrained MAPLE model, a MAPLE instance was trained from scratch using the combined SRRSH dataset (SRRSH-24 and SRRSH-141, 285 samples total) with a 5-fold cross-validation. Its performance was evaluated on the SRRSH-141 samples in each testing split to ensure comparability with other DF-IM-TL results. Training used the 70,110 MAPLE CpGs and default MAPLE neural network layer configuration, with encoder regularization on, learning rate of 0.0001, batch size of 256 and 6000 epochs.

### Clinical clock training

Among 59 collected clinical biomarkers, 20% of features with least correlation with CA was screened out. Missing values were filled (filling 0, all features have missing ratio < 10%). Data was standardized before feeding into an elastic net model for training via 5-fold cross-validation. The hyperparameters (alpha and l1_ratio) were optimized by an independent 5-fold cross-validation.

### JSD calculation and ALS classification

Jensen-Shannon Divergence (JSD) was calculated between estimated (by Gaussian kernel density estimation) probability density function of delta-age distributions predicted by each clock individually. The bandwidth parameter in KDE is automatically determined using the “scott” method. ALS classification was performed via support vector machine (SVM) classifier with a linear kernel using the per-clock delta-ages. Classification performances were evaluated using leave-one-pair-out cross-validation.

## Results

### Discordance of epigenetic profiling across array and HTS platforms

We compared four profiling technologies, Illumina MSA and EPICv2 (arrays) alongside iGeneTech Galaxy and Twist (HTS), using the SRRSH-24 cohort (24 participants; six datasets spanning both gDNA and cfDNA sources with technical replicates; Fig. 1a). The remaining 141 participants (SRRSH-141) were reserved for independent validation. The above methods were selected to represent both established and emerging methylation profiling technologies (Methods).

**Fig. 1.**
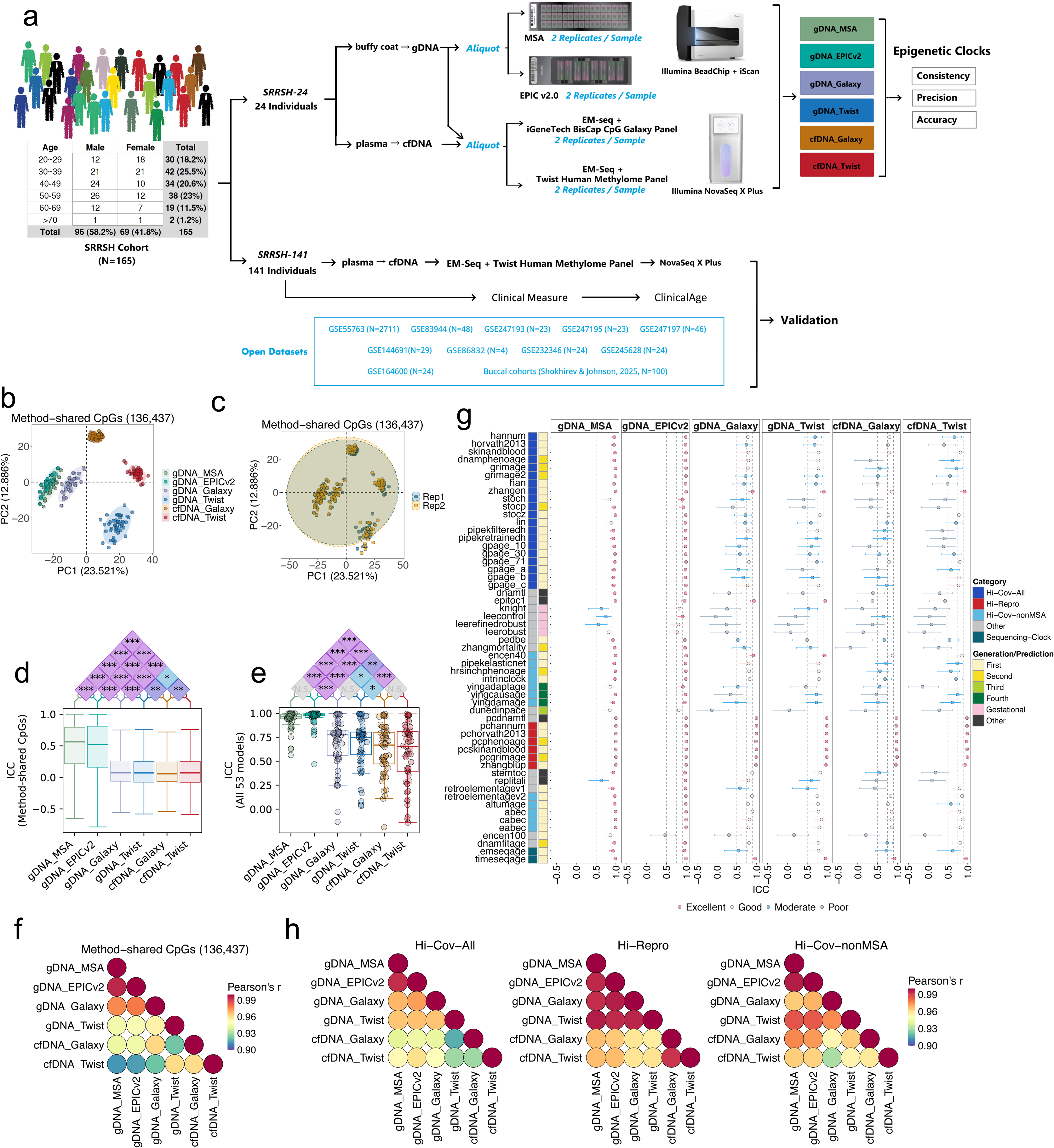
Study design and evaluation of methylation data reproducibility and consistency across six SRRSH-24 datasets. **a**, The overall design of the study. **b-c**, PCA of all samples colored by datasets (**b**) or replicates (**c**), using beta-values of Method-shared CpGs (N = 136,437). **d**, Intraclass Correlation Coefficients (ICCs) of Method-shared CpGs across six datasets. **e**, Comparisons of per clock ICCs of all array-trained clocks (N = 53). **f**, Pearson correlations among datasets computed using the mean beta-values of Method-shared CpGs. **g**, ICCs of 55 epigenetic (53 array trained and 2 HTS-trained) clocks benchmarked in the study. Data are presented as ICC estimates with 95 % confidence intervals and colored by grouped ICC values as follows: Excellent: 0.9 ≤ ICC, Good: 0.75 ≤ ICC < 0.9, Moderate: 0.5 ≤ ICC < 0.75, Poor: ICC < 0.5. Clocks were classified according to two different rationales: (1) By generations: First — Chronological age prediction; Second — Phenotypic age or mortality prediction; Third — Pace of aging prediction; Fourth — Causal clocks; Gestational — Gestational age prediction; Other: Telomere length or mitotic age prediction. (2) By CpG coverage across the methylation profiling methods. **h**, Pearson correlations across six datasets calculated using the mean of clock predictions for three clock performance categories. P-values in (**d**) and (**e**) were calculated by two-tailed block permutation test with 10,000 permutations, then adjusted for multiple comparisons using the Benjamini-Hochberg procedure. ***, p < 0.001; **, p < 0.01; *, p < 0.05; N.S., not significant. The center line of each boxplot marks the median, the box body marks the 1^st^ and 3^rd^ quartiles, the caps mark the min and max values.

### Coverage-induced discordance between array and HTS platforms

Although HTS capture panels offer significantly larger total marker sets (Galaxy: ∼2.0M; Twist: ∼3.98M) than arrays (MSA: ∼270K; EPICv2: ∼935K) (Fig. S1a), this genomic breadth did not translate to higher clock-specific coverage (Fig. S1b). Consequently, this disparity causes individual clocks on average to miss 30% biomarkers when applied to HTS data. In addition, “coverage drift” occurs in HTS platforms where observed CpG coverage exceeds specifications (Galaxy: +4.8%; Twist: +1.06%). These flanking region capture, probe redundancy, or broader coordinate overlap can further reduce biomarker fidelity. Overall, the Twist panel achieved the highest overall feature retention among these clocks, due to a more faithful site preservation of the legacy methylation array (e.g., 450K) that constitute the training foundation for the majority of established epigenetic clocks.

### CpG-level reproducibility and technical noise

Seamless cross-platform deployment of epigenetic aging biomarkers requires strict consistency (agreement across other methods) and reproducibility (agreement between technical replicates). However, benchmarking 136,437 shared genomic sites revealed a profound systematic bias that separates HTS from arrays (Fig. 1b-c). HTS platforms exhibited significantly weaker reliability at individual biomarker loci (P < 1×10^-5^; Fig. 1d), primarily due to a massive enrichment of unreliable, extreme methylation values (0 or 1) that comprised 60.5-79.9% of HTS data compared to just 30.8-35.7% in arrays (Fig. S2). Notably, such biomarker observations are often considered unreliable for age predictions [7,20].

This analytical noise directly undermines the clinical utility of biological age prediction models, severely depressing predictive reproducibility in HTS environments (ICC = -0.188-0.997, median 0.721) relative to standard arrays (ICC = 0.469-0.999, median 0.950; Fig. 1e). This structural divergence hinders the direct transferability of legacy age-prediction biomarkers; while biomarker consistency between standard array platforms approaches parity (Pearson’s R = 0.9947), it falters significantly across all HTS-involved comparisons (R = 0.9125-0.9781; Fig. 1f).

### Clock categorization by cross-platform robustness

To facilitate clinical translation, we categorized the 53 evaluated epigenetic clocks based on their cross-platform robustness (Fig. 1g, Table S1). Six 1^st^-generation clocks were designated as “Hi-Repro” due to their exceptional test-retest stability (ICC ≥ 0.9; median replicate difference (RD) 0.17-3.81; Fig. S3, S4). These robust models naturally filter out technical noise during individual age prediction, although without guaranteed accuracy at cohort level. Hi-Repro clocks demonstrated the highest cross-platform consistency in gDNA, while still suffered performance declines in cfDNA (Fig. 1h, S5). Notably, only 17 of the evaluated models (including all Hi-Repro clocks) successfully maintained high BA prediction consistency across all platforms (Fig. S5). We also identified 32 additional models that retained sufficient CpG coverage (≥ 80%) across platforms, classifying them as high-coverage candidates (“Hi-Cov-All” and “Hi-Cov-nonMSA”) for cross-platform prediction. Ultimately, these highly reproducible and high-coverage models emerge as the most viable epigenetic biomarkers for reliable, cross-platform aging assessments.

### Impact of feature selection and platform divergence on predictive stability

To evaluate whether the inherent feature selection that occurs during the clock training naturally filters out technically unstable biomarkers, we benchmarked 2,727 “Clock-shared CpGs” shared by Hi-Cov-All and Hi-Repro clocks. We hypothesized that the rigorous selection of aging relevant loci might inherently bypass platform noise. Although this curated subset of biomarkers yielded a marginal improvement in global consistency (mean Pearson’s R = 0.97; Fig. S1h) compared to the unselected background (R = 0.95; Fig. 1f), substantial platform-specific stochasticity persisted (Fig. S1e-f). Array-based platforms continued to deliver superior predictive consistency (Fig. S1c-d) and reproducibility (Fig. S1g) over HTS platforms. This demonstrates that while the original model training effectively isolates aging signals, this biological feature selection alone is insufficient to filter out platform-based technical noise. Consequently, deploying these legacy clocks for HTS-based age prediction mandates dedicated downstream computational adaptation.

### BA prediction accuracy and the penalty of platform dependency

Evaluating epigenetic clocks in the SRRSH-24 cohort revealed a significant BA prediction accuracy penalty when applying to HTS-based cfDNA. Compared to array-based gDNA (mean absolute error [MAE] = 2.03-23.08 years, median 7.39; R = 0.44-0.96, median 0.80; Fig. 2a-c, S6) as a clinical reference, applying legacy models directly to cfDNA yielded elevated biological age prediction errors (MAE = 5.26-18.75 years, median 10.83) and blunted correlations with chronological age (R = 0.40-0.92, median 0.76).

**Fig. 2.**
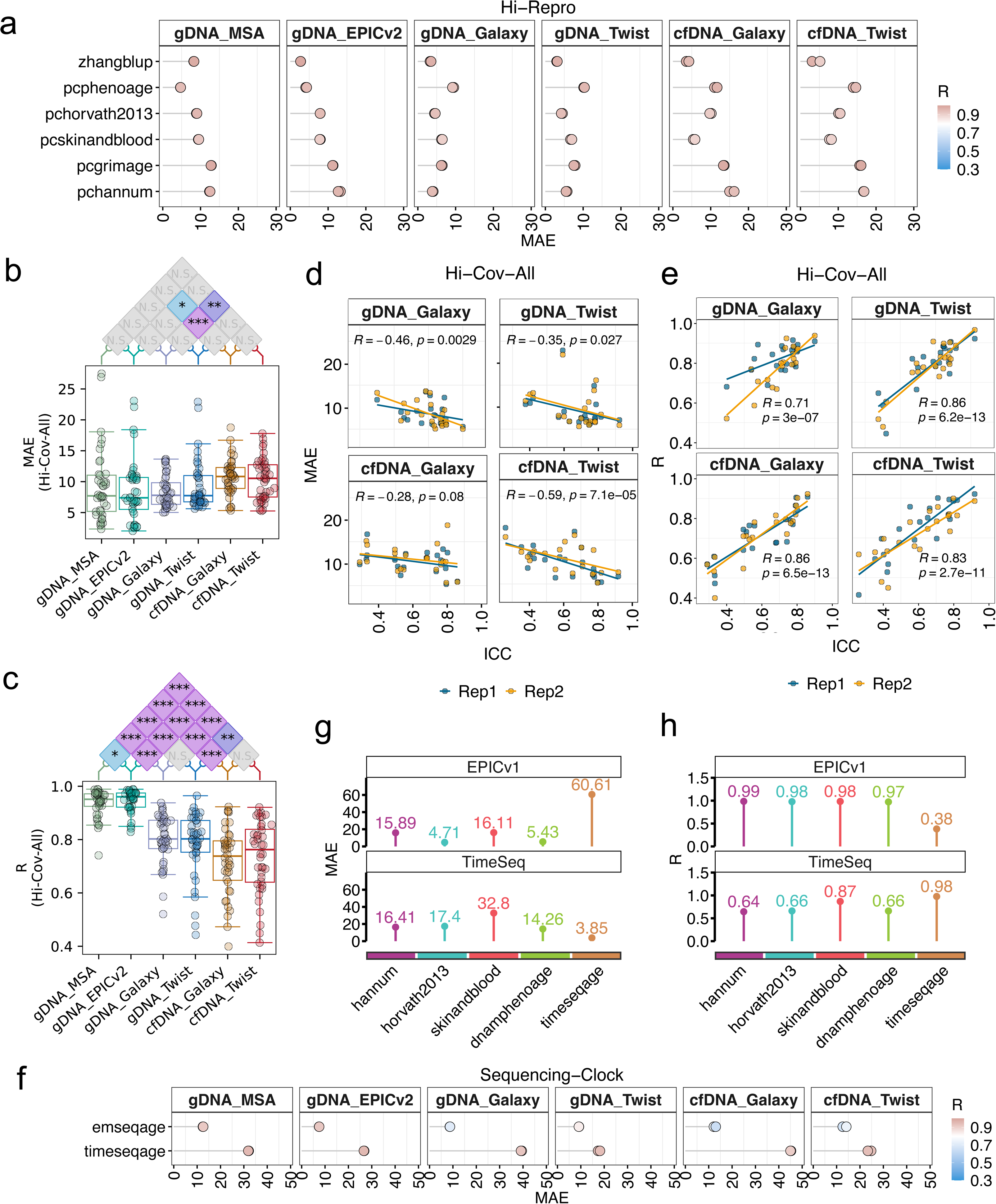
Decline of epigenetic clock performance in 6 SRRSH-24 HTS-based datasets. **a**, Lollipop plot showing Pearson’s R and MAE for 6 Hi-Repro clocks. Each dot represents a replicate. **b-c**, Boxplots showing comparisons and statistical significance of (**b**) MAE and (**c**) Pearson’s R for Hi-Cov-All clocks. The center line of each boxplot marks the median, the box body marks the 1^st^ and 3^rd^ quartiles, and the caps mark the min and max of RD values respectively. **d**, Scatter plot showing the correlation between clock ICC and MAE in the four SRRSH-24 HTS datasets. **e**, Scatter plot showing the correlation between clock ICC and R in the four SRRSH-24 HTS datasets. Lines represent linear regression fits for each replicate group respectively. **f**, Lollipop plot showing Pearson’s R and MAE for the 2 HTS-trained clocks. Each dot represents a replicate. **g-h**, Performance evaluation of 4 array-trained clocks (hannum, horvath2013, skinandblood and dnamphenoage) and an HTS data-trained clock (timeseqage) cross-applied to the array-based dataset (GSE245628, labelled as EPICv1) and HTS-based dataset (GSE232346, labelled as TimeSeq). Only the 24 shared samples in both GSE245628 and GSE232346 were analyzed. The p-values in (**b**) and (**c**) were calculated by two-tailed block permutation test with 10,000 permutations, then adjusted for multiple comparisons using the Benjamini-Hochberg procedure. ***, p < 0.001; **, p < 0.01; *, p < 0.05; N.S., not significant; The R and p-values in (**d**) and (**e**) were calculated by Pearson correlation at the replicate level.

Crucially, test-retest reliability (ICC) strongly correlated with overall diagnostic accuracy, confirming that HTS noise directly compromises biomarker validity (Fig. 2d-e). While highly resilient models like zhangblup maintained strong cross-platform accuracy (MAE ≤ 5.2 years), their clinical utility still deteriorated under critical feature missingness (e.g. 75% missing, Fig. 2a, S1b, S7). These observations confirm that most legacy clocks are not tuned to the biomarker stochasticity and fidelity in HTS-based cfDNA methylation data, and the induced model behavior hampers their application for cfDNA-based BA assessments.

Furthermore, performance profiling demonstrated a profound platform dependence. Epigenetic models exhibited distinct “platform lock-in”: array-trained models (e.g., hannum) vastly outperformed sequencing-trained models (timeseqage) on array data, and vice versa on sequencing datasets (Fig. 2f-h). These results indicate that clocks are inherently tuned to dataset-or platform-specific patterns, necessitating targeted recalibration before clinical cross-platform deployment. In this study, further benchmarking and adaptation focused on array-trained clocks because of their better standardized feature set and more versatile clinical applications beyond CA predictions.

### Optimizing sequencing depth for HTS-based epigenetic clock applications

To determine the mechanisms driving diminished diagnostic reliability in HTS-based cfDNA data, we isolated sequencing depth as a technical confounder and systematically evaluated its effect on biomarker stability. Its influence on methylation signal fidelity was assessed by modeling the stochasticity of observed read counts and the inherent analytical errors in ratio approximation. Both simulation-based and downsampling-based benchmarking on the SRRSH-141 cohort demonstrated that MAE and RD stabilize at a minimum mean target depth of 10×, with diminishing returns observed beyond 20× for both tested linear and non-linear clocks (Fig. 3a-d, S8). Interestingly, these thresholds coincide with standards for whole genome bisulfite sequencing [35,36] discovered from distinct rationales. Notably, all HTS datasets generated in the present study satisfied this 10× minimum requirement (Fig. S9).

**Fig. 3.**
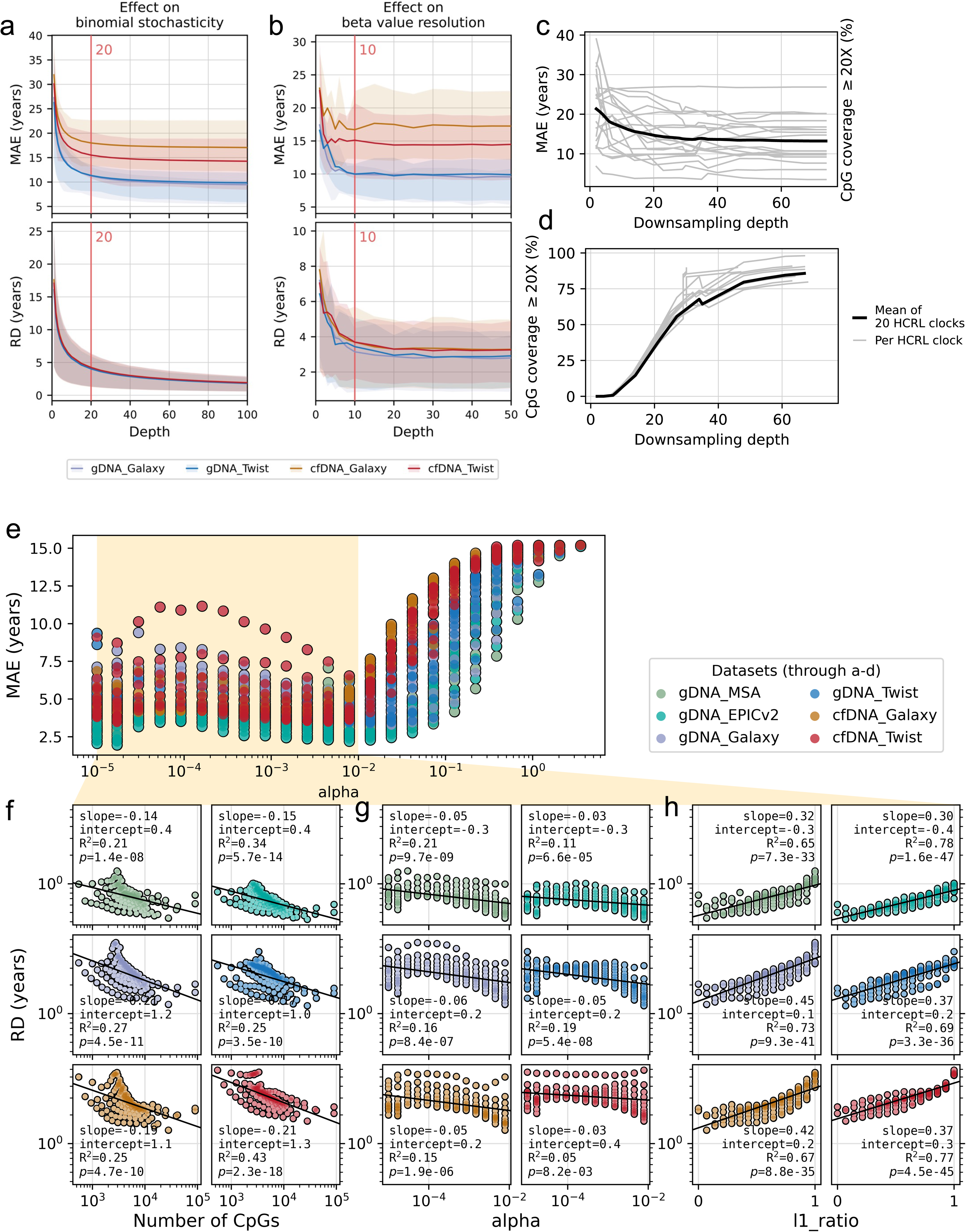
Optimization of epigenetic clock applications for HTS-based methylation data via sequencing depth and model selection. **a**, Investigation of the sequencing depth threshold via beta-value stochasticity effect at individual CpG level on HCRL clocks. **b**, Investigation of sequencing depth threshold via beta-value approximation precision on HCRL clocks. **c-d**, Investigation of the sequencing depth requirements with downsampling simulations, using five samples from the SRRSH cohort with raw sequencing output ≥ 50Gb. **c**, Age prediction MAE at different downsampling depths. **d**, The increase in the fraction of CpGs with ≥ 20× coverage slowed down beyond the 20× downsampling depth. **e**, A series of elastic nets were trained on the dataset GSE55763 (N = 2711) and tested with the 6 datasets from this study. **f**, The negative correlation observed between the number of CpGs and age prediction RD. **g-h**, Comparison of correlation between hyperparameter (alpha and l1_ratio) and RD. R^2^ and p-values were calculated by Pearson correlation.

### Feature density and L2 regularization enhance reproducibility

To delineate the primary architectural drivers of biomarker reliability, we selected 20 High Coverage/Reproducibility Linear (HCRL) clocks from the Hi-Cov and High-Repro categories, then analyzed them across 11 datasets (Table 1, Fig. S10). A strong negative correlation was observed between number of CpGs and RD, indicating that high-density feature sets significantly fortify models against technical variance. These results reinforce prior evidence that broader genomic including provides a structural buffer against platform artifacts [7,37], suggesting that the physiological aging signal is widely distributed across the epigenome.

**Table 1.** Linear regression results between the age prediction replicate difference (RD) and the number of CpGs in 20 HCRL clocks. HCRL denotes 20 linear-architecture clocks classified as Hi-Cov-All or High-Repro. Negative correlations indicate larger number of CpGs is associated with lower age prediction discrepancies between technical replicates. P-values were adjusted for multiple testing using the Benjamini-Hochberg procedure. Full regression outputs for each dataset are provided in Fig. S10.

| **Dataset** | **Slope** | **Intercept** | **Pearson's R** | **p** |
| --- | --- | --- | --- | --- |
| gDNA_MSA | -0.36 | 2.34 | -0.94 | **3.65E-10** |
| gDNA_EPICv2 | -0.24 | 1.30 | -0.87 | **4.67E-07** |
| gDNA_Twist | -0.28 | 2.87 | -0.90 | **4.91E-08** |
| gDNA_Galaxy | -0.31 | 3.01 | -0.93 | **3.49E-09** |
| cfDNA_Twist | -0.29 | 3.04 | -0.91 | **2.69E-08** |
| cfDNA_Galaxy | -0.28 | 3.01 | -0.90 | **5.86E-08** |
| GSE83944 | -0.18 | 0.75 | -0.83 | **7.78E-05** |
| GSE247193 | -0.19 | 1.60 | -0.84 | **5.65E-05** |
| GSE247195 | -0.22 | 1.14 | -0.85 | **2.99E-05** |
| GSE247197 | -0.14 | 0.55 | -0.67 | **4.62E-03** |
| GSE55763 | -0.18 | 0.93 | -0.81 | **1.65E-04** |

To isolate the mechanisms underlying this stability, we systematically tuned the elastic net hyperparameters: regularization strength (alpha) and the L1/L2 mixing parameter (l1_ratio). The correlation between feature count and RD was reaffirmed within the optimal alpha range (≤ 0.01, Fig. 3e-f). Sensitivity analyses revealed that l1_ratio serves as a profoundly more potent determinant of reproducibility (R^2^ = 0.67-0.78; p = 1.6×10^-47^ - 7.3×10^-33^) than alpha (R^2^ = 0.05-0.21; p = 9.7×10^-9^ - 8.2×10^-3^; Fig. 3g-h). Overall, biasing the model toward an L2 (Ridge) penalty preserves a broad distribution of small-effect epigenomic aging signals, which effectively increased age prediction reproducibility. Consequently, L2-heavy architectures are essential for achieving reproducible predictions in sequencing-based applications.

### Clock accuracy improvement with the DF-IM-TL adaptation pipeline

To mitigate aging signal artifacts in HTS data, we developed a three-stage HTS data processing pipeline: depth filtering (DF), imputation (IM), and transfer learning (TL).

#### Stochastic noise mitigation via DF and IM

The initial DF stage systematically excludes genomic loci with sub-optimal sequencing coverage, flagging them for subsequent processing by IM. Evaluation across eight targeted sequencing cohorts revealed a high prevalence of artifacts manifesting at absolute distribution extremes (beta-values of 0 or 1). These artifacts accounted for 18.4% to 61.3% of the total assessed CpGs and compromised 2.0% to 34.7% of the critical loci utilized by robust clinical epigenetic models (Fig. 4a, S11-S12, Table S3). Deeper analyses confirmed that these extreme beta-values represent technical noise rather than genuine epigenetic states, evidenced by their distinct depth profiles compared with biologically plausible ones (Fig. S13-S14).

**Fig. 4.**
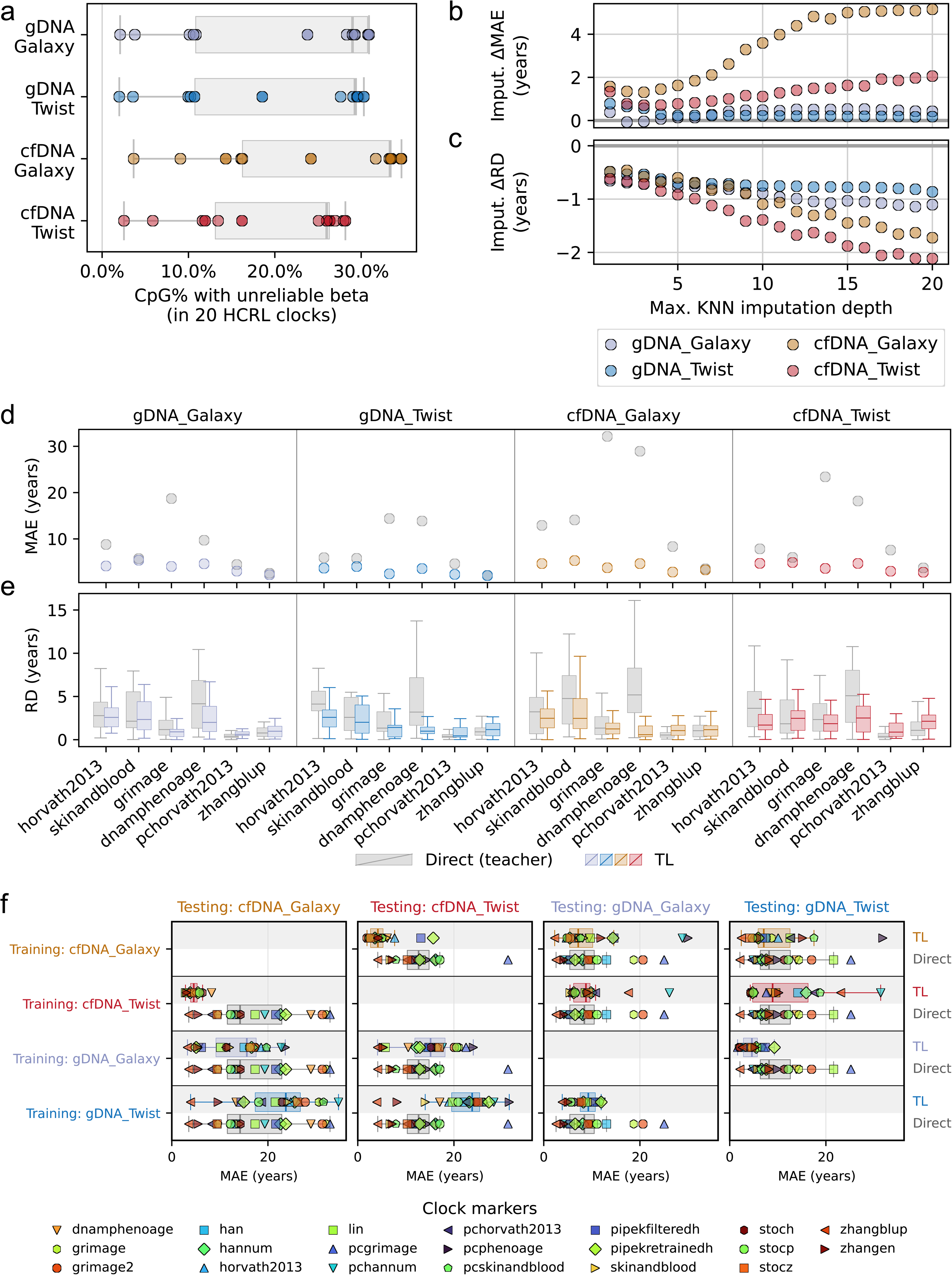
Investigating depth filtering (DF), imputation (IM) and transfer learning (TL) to improve epigenetic clock performance across array and HTS platforms. **a**, HCRL clocks are prone to unreliable beta-values originating from HTS-based methylation profiling technologies (Details are shown in Table S3). **b-c**, The K-nearest-neighbor (KNN) imputation technique was tested at varying depth cut-offs; only CpG sites below the cut-off were imputed. **d-e**, Comparison of age prediction MAE (**d**) and RD (**e**) between teacher and student (distillation) clocks, assessed with 5-fold cross validation. **f**, Generalizability benchmarking of transfer learning across HTS capture panels and DNA types. The center line of each boxplot marks the median, the box body marks the 1^st^ and 3^rd^ quartiles, and the caps mark the min and max values.

The imputation phase then is designed to rescue overall methylation profiling reliability. K-nearest neighbor (KNN) imputation consistently tightened test-retest variance across all platforms, reducing RD by 0.7-2.1 years in cfDNA and 0.7-1.1 years in gDNA (Fig. 4b). Other alternative imputation strategies such as mean, median imputation offered comparable reproducibility improvements and accuracy trade-offs with a notable exception for 0-filling (Fig. S15). This is potentially because basic heuristic approaches risk amplifying the stochastic noise inherent to the profound cellular heterogeneity of cfDNA. Notably, zhangblup demonstrated exceptional clinical resilience, showing consistent performance enhancement across all datasets (Fig. S16).

#### TL for cross-platform alignment

To harmonize legacy array-derived epigenetic biomarkers with HTS data, we incorporated model distillation (Teacher-Student transfer learning). This platform-agnostic strategy leverages an established, clinically validated array model to carefully calibrate an HTS counterpart. To reduce overfitting, we incorporated PCA-based dimensionality reduction (Methods) and five-fold cross-validation was incorporated. Distillation significantly improved prediction accuracy and reproducibility across six representative first- and second-generation clinical epigenetic clocks (Fig. 4d-e). Furthermore, TL effectively preserved the utility of noise-susceptible models such as dnamphenoage. Crucially, this recalibration selectively neutralized platform-induced artifacts while retaining high sensitivity to DNA origin; performance gains were optimal when adaptation and targeting datasets shared the same DNA origin (Fig. 4f).

### Age prediction benchmark of the DF-IM-TL pipeline

The DF-IM-TL pipeline was benchmarked in two independent datasets. In the Buccal cohort (N=100), TL-inclusive strategies universally outperformed others, with the TL-only strategy performing best, yielding MAE reductions of 1.5 - 42.0 (median 10.3) years and CA correlation improvement of 0.02 - 0.78 (median 0.15) (Fig. 5a-b, S17). Similarly, in the SRRSH-141 cohort, the IM+TL strategy achieved the highest reduction of MAE by 0.89 - 24.0 (median 7.7) years and improving Pearson’s R by 0.01 - 0.78 (median 0.18) (Fig. 5c, S18-S19). Notably, independent tests show that the DF-IM-TL pipeline also outperformed the existing MAPLE framework in MAE while maintaining comparable reproducibility (Fig. 5a, S18), regardless of whether MAPLE was used with its pre-trained model or trained from scratch using default parameters (Methods). MAPLE is specifically designed to handle cross-platform biases [38]. The DF-IM-TL pipeline also outperformed three other commonly used adaptation algorithms—quantile mapping, ComBat, and correlation alignment (CORAL)—each attempting to align HTS datasets to gDNA_EPICv2 before prediction with original HCRL clocks. The results in Fig. 5d suggest DF-IM-TL’s highly specialized efficacy in correcting HTS platform biases in epigenetic age prediction.

**Fig. 5.**
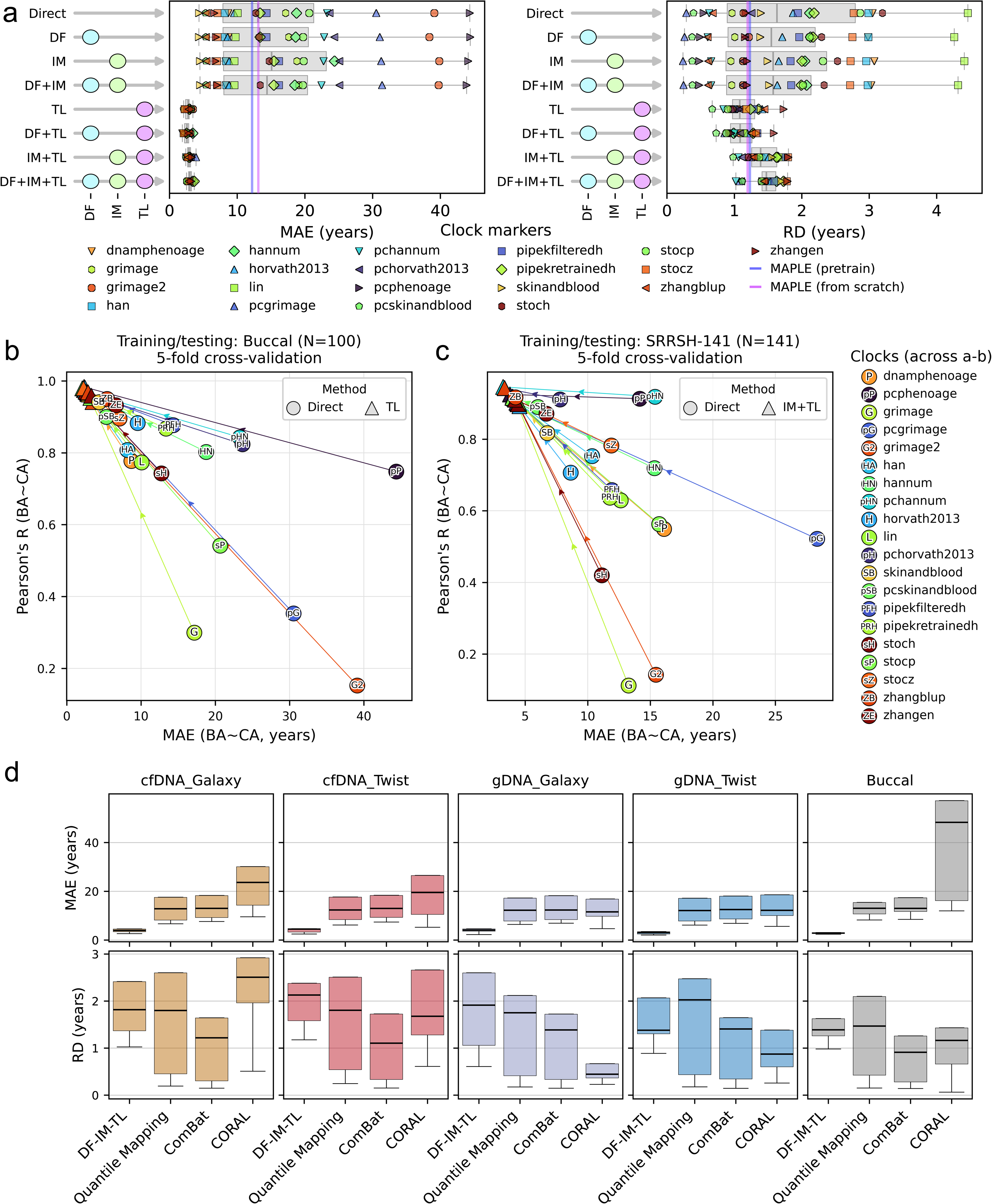
Testing and validating the DF-IM-TL epigenetic clock adaptation pipeline for biological age prediction on HTS platforms. **a**, Evaluation of all DF-IM-TL combinations for 20 HCRL clocks on the Buccal cohort (N = 100). **b**, Performance improvements via the best identified strategy (TL-only) with respect to CA on the Buccal cohort. **c**, Performance improvements via the best identified strategy (IM-TL) with respect to CA on the SRRSH-141 cohort (N = 141). **d**, Adaptation efficacy comparison between DF-IM-TL and three other commonly used adaptation strategies.

### Constructing the clinical clock

To test the DF-IM-TL pipeline’s ability to improve concordance between HCRL clocks and clinically informed reference, we utilized a clinical data-derived BA as a reference. The ClinicalAge clock was constructed using 59 biomarkers from the combined SRRSH-24 and SRRSH-141 cohort (N=189), achieving a MAE of 6.48 years against CA (Fig. 6a). Analysis of the top 10 features revealed strong weightings toward metabolic, hepatic, and inflammatory markers. Specifically, high positive correlations for glycated hemoglobin and blood glucose reflect age-related glucose homeostasis decline. Additionally, positive coefficients for aspartate aminotransferase (AST) and alkaline phosphatase (ALP) reflect age-related cellular damage and liver stress. In contrast, alanine transaminase (ALT) exhibited a strongly negative coefficient; this suggests the model captures the well-established AST/ALT ratio, a known marker of liver aging, rather than treating these biomarkers independently. These key features strongly align with established clinical clocks like PhenoAge [5,39] and GOLD BioAge (Fig. 6b). It is worth noting that we explicitly trained this clock from scratch since existing clinical models (e.g. PhenoAge) often incorporates CA in the formula, contrasting with the preference of using a CA-independent metric as age acceleration (AA, the discrepancy between BA and CA) reference.

**Fig. 6.**
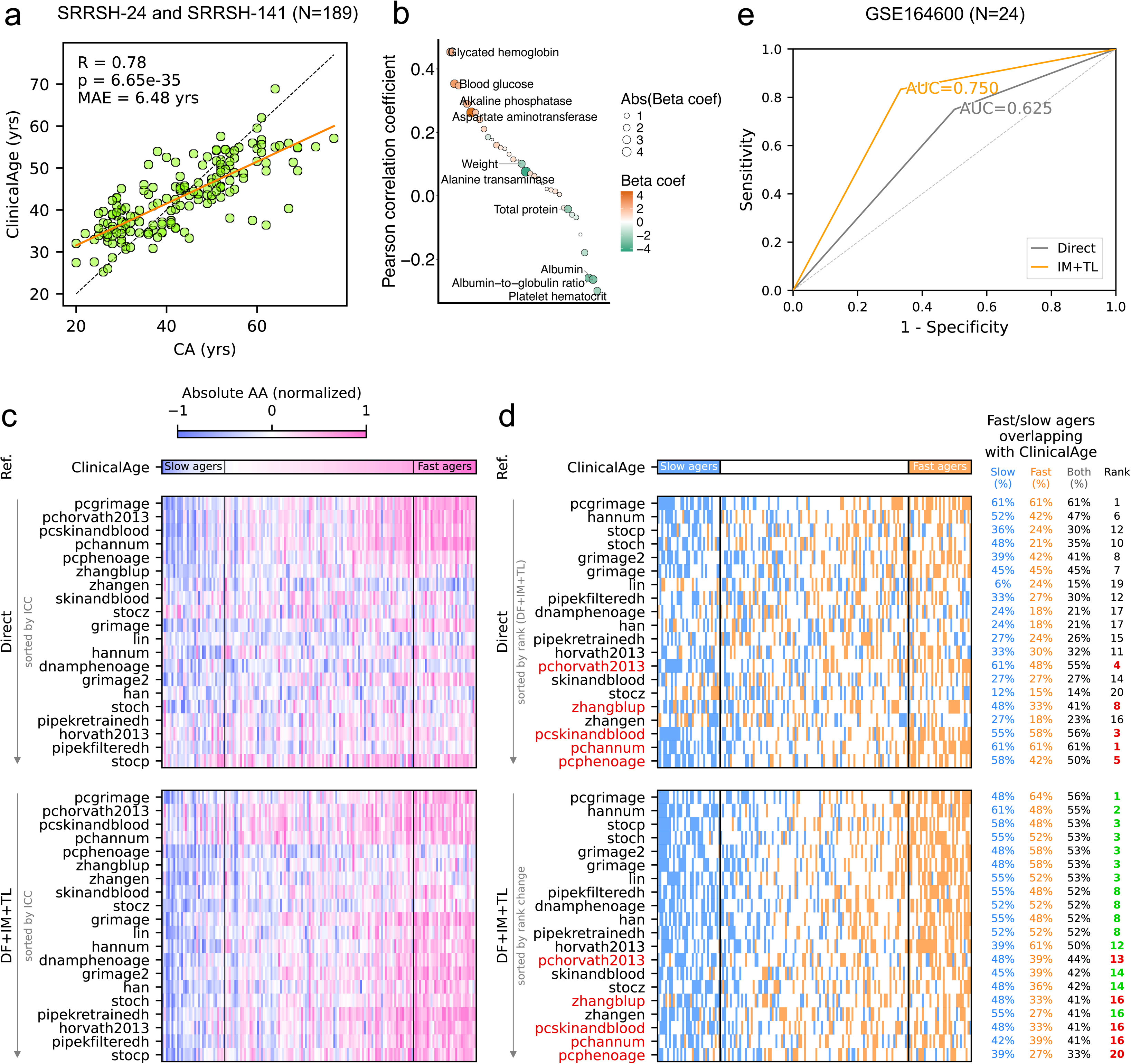
Testing and validating the DF-IM-TL epigenetic clock adaptation strategies using clinical data. **a-d**, TL improved identification of clinical fast agers using legacy epigenetic clocks. **a,** MAE and R of an elastic net clinical clock trained from collected clinical biomarkers to predict CA. **b,** Feature importance analysis of clinical clock. **c,** Concordance of normalized absolute Age Acceleration (AA) between prediction by legacy clock (Direct), prediction after DF-IM-TL, and clinical clock as reference. Clocks are ranked in descending order of prediction ICCs. **d,** Concordance between fast ager and slow ager identifications between prediction by legacy clock (Direct), prediction after DF-IM-TL, and clinical clock as reference. Deteriorations are marked red in DF-IM-TL result with respect to Direct, and improvements are in green (rank only). **e-f,** Validation by disease classification performed on a public dataset GSE164600 (N = 24) with healthy and amyotrophic lateral sclerosis (ALS) labels. **e**, Comparison of Jensen-Shannon Divergence (JSD) to evaluate the distributional distances between ALS and normal samples before and after DF-IM-TL. **f**, Maximum AUC achieved using as few as 4 top ΔJSD clocks, surpassing the non-adapted prediction by 0.125 AUC.

### DF-IM-TL improves identification of fast agers

Fast ager identification is pivotal for aging clocks preceding aging interventions. Following DF-IM-TL adaptation, HCRL clocks demonstrated significantly improved AA concordance with ClinicalAge (Fig. 6c). Biologically, this indicates that the pipeline successfully filters HTS noise to recover true physiological aging signals. This recovery of biological signal was particularly profound for noise-susceptible, low-ICC clocks, whereas inherently stable (Hi-Repro) models showed marginal changes. By defining the top and bottom 20% of ClinicalAge AA as fast and slow agers respectively, we confirmed that the pipeline significantly enhances the clinical accuracy of these categorizations across legacy models (Fig. 6d). Therefore, DF-IM-TL serves as a practical tool to enhance the ability of recovering meaningful aging trajectories from noisy HTS methylation profiles via legacy clocks, though it may be bypassed for clocks with high baseline reproducibility.

### DF-IM-TL improved AA-based disease detection

We tested the DF-IM-TL framework using an age-balanced amyotrophic lateral sclerosis (ALS) cfDNA dataset (N=24; GSE164600), as a proof-of-concept evaluation of disease sensitivity. Although chronological age is the primary risk factor for ALS, AA serves as a more discriminative biomarker for capturing heterogeneous clinical decline [40,41]. After adaptation, the Jensen-Shannon Divergence (JSD) between ALS and control AA distributions widened significantly (Fig. S20). This further increase AA-based disease classification AUC by 0.125 as tested by an SVM classifier (Fig. 6e). These findings confirm that by filtering HTS technical noise to recover genuine pathological signatures.

## Discussion

Integrating array-trained epigenetic clocks with HTS-based cfDNA data demands rigorous quantitative reconciliation of platform-specific biases. Addressing the scarcity of cross-platform benchmarks, the SRRSH-24 dataset provides a comprehensive technical replicate matrix evaluating both DNA substrates and profiling technologies. This benchmarking exposes a fundamental diagnostic challenge: HTS methodologies introduce severe technical stochasticity, significantly degrading individual biomarker (CpG-level) reproducibility compared to standard arrays. Because technical reproducibility is necessary but insufficient to guarantee the preservation of biological signals, future HTS-adapted biomarkers must prioritize reproducibility, measured via RD, alongside traditional accuracy metrics like MAE. This dual-metric approach is essential to successfully decouple genuine physiological aging from technical assay noise.

Adapting epigenetic clocks for HTS data requires intrinsic resilience against platform-specific bias and noise. Our results show that this robustness can be achieved through collective stability: utilizing high-density CpG sets to distribute small-effect signals across the epigenome, buffering age predictions against site-specific stochasticity. For linear models, this necessitates shifting biomarker selection away from sparsity-oriented L1 regularization toward an L2-heavy elastic net architecture. By prioritizing L2 penalties, this framework undergoes a similar mathematical structure associated with the high cross-platform reproducibility of PCA- and BLUP-based models (e.g., zhangblup; Fig. S3, S4a; see Supplementary Discussion for their relationships). Consequently, integrating an L2-heavy elastic net during biomarker selection is highly recommended as a foundational design principle for adapting legacy linear epigenetic clocks, ensuring highly reliable age prediction in HTS-based clinical applications.

To neutralize platform-induced artifacts beyond the use of high-density biomarker sets, the sequential integration of Depth Filtering (DF), Imputation (IM), and Transfer Learning (TL) provides a robust adaptation framework. This pipeline progressively isolates the physiological aging signature through three targeted phases: (1) DF restricts individual CpG-level stochasticity by excluding sub-optimal genomic loci; (2) IM systematically mitigates residual technical artifacts to stabilize predictive variance; and (3) TL recovers the genuine biological signal for age prediction. The synergistic application of these three stages maximizes diagnostic accuracy and successfully harmonizes predictive discordance across sequencing platforms, which is critical for cross-sectional comparison between biomarkers. Ultimately, this methodology drives highly efficient “technological decoupling”—the systematic unmasking of true epigenetic aging signals from the inherent stochastic nature of HTS.

Beyond this specific adaptation pipeline, these methodologies offer broad utility for the clinical deployment of epigenetic biomarkers. **DF** guideline also establishes a recommended sequencing depth threshold, functioning as a vital quality-control standard to guarantee the diagnostic reliability of BA estimations in clinical settings. **IM** also revealed the profound impact of cellular heterogeneity on cfDNA-based aging assessments. The cfDNA biomarkers exhibited a more pronounced accuracy trade-off than standard genomic DNA, likely driven by the complex tissue-of-origin heterogeneity inherent to cfDNA [19], which can constrain the clinical generalizability of pan-tissue aging biomarkers [15] and skew clinical predictions when relying on basic heuristic imputation algorithms. Integrating cell-type deconvolution prior to imputation may resolve this limitation by incorporating biologically informed prior knowledge [19,42]. **TL** via model distillation acts as a critical mechanism for recovering the underlying physiological aging signal and optimizing predictive accuracy. Because this distillation framework is fundamentally model-agnostic, it offers universal applicability, enabling the seamless translation of a broad spectrum of legacy aging biomarkers into modern, actionable clinical diagnostics.

Overall, by bridging the data-level divide between legacy array-based knowledge and modern sequencing technologies, the DF-IM-TL framework methodology effectively recovers methylation profiles previously considered lost to technical noise. Consequently, by restoring the clinical validity of noise-susceptible legacy clocks, this integrated pipeline establishes a highly reliable analytical foundation for the non-invasive, longitudinal monitoring of physiological health.

While this study establishes a robust methodological foundation, certain limitations remain. Due to the limited sample size and lack of data, no sex- or ethnicity/race-related effects were evaluated. Furthermore, all datasets were cross-sectional without longitudinal and intervention-oriented analysis, precluding insights into temporal methylation dynamics. These limitations may restrict the generalizability of our findings to broader populations and dynamic aging processes. Future work should extend to longitudinal cohorts to evaluate how adaptation influences sensitivity to age-related interventions. While linear models were prioritized for their interpretability and high performance in high-dimensional spaces, the true biological relationship between methylation and aging is not necessarily linear. Therefore, though linear epigenetic clocks can already achieve high accuracy [6,7,43], future research into advanced clocks and domain adaptation methods may further improve cross-platform accuracy [25,44]. Additionally, the challenge of tissue-of-origin heterogeneity in cfDNA remains a critical variable that warrants further investigation to optimize imputation and recalibration strategies.

## Conclusion

This study provides a comprehensive methodological framework for the cross-platform application of epigenetic clocks. By contributing the SRRSH-24 benchmarking resource, formalizing depth requirements, and validating the DF-IM-TL pipeline, we offer a clear path towards HTS-based aging research. The proposed pipeline framework bridges the extensive gallery of array-derived epigenetic clocks to be reliably deployed in the emerging HTS-based epigenetic aging and liquid biopsy diagnostics.

## Acknowledgement

Special thanks to the high-performance computing cluster provided by Liangzhu Laboratory, Zhejiang University for computation support.

## Permission statement

The authors declare no extra permission statements.

## Authors’ contributions

C.O. and G.Z. conceived and supervised the study. G.L., C.O., and G.Z. managed the overall project. Cohort management: L.C., X.C., Z.Y., D.T., X.F., X.H., and V.T. organized the cohort, collected human blood samples, and coordinated resources. X.H. and V.T. also provided essential medical and technical support. Data generation: C.O., Y.G., S.J., B.H., and C.T. performed sample collection, DNA extraction, and library preparation. Bioinformatics analysis: J.W. and W.H. developed the bioinformatics pipeline. J.W. preprocessed HTS and methylation array data and performed downsampling analysis. W.H. conducted reproducibility, consistency, and accuracy analyses. G.L. investigated clock complexity, unreliable beta filtering, transfer learning, and combination strategies. X.Z. contributed to statistical and machine learning analyses, with advisory support from Y.W. Manuscript preparation: G.L., W.H., X.Z., J.W., and C.O. wrote the manuscript. All authors reviewed and approved the final manuscript.

## Data availability

The de-identified data generated in this study, including processed methylation matrices, sequencing depth matrices and non-sensitive subject metadata are available at zenodo.org under accession 17453844 (DOI: 10.5281/zenodo.17453844). Raw data available at GSA for Human (https://ngdc.cncb.ac.cn/gsa-human/) under accession HRA014554.

## Code availability

The code related to this project is publicly accessible at https://github.com/regen-bio/cfDNA-HTS-Clock-Analysis.git.

