## Supplementary Information for "A robust and integrated framework for cross-platform adaptation of epigenetic clocks in cell-free DNA sequencing"

### Supplementary Discussion

#### ZhangBLUP is mathematically related to ridge regression

Since ZhangBLUP, PCA-preprocessed models, and ridge regression all showed a similar effect in reducing RD, we further interrogated their algorithm details and revealed that these models are indeed mathematically related rather than being coincidental to have this RD-reducing effect. Best linear unbiased predictor (BLUP) is a special case of linear mixed model (LMM):

$$y=X\beta+Zu+\epsilon$$

Where:

X is the fixed coefficient matrix.

$\beta$ is the fixed effect size vector.

Z stands for the random effect coefficient matrix.

$u\sim N(0,\sigma_{u}^{2}I)$ is the random effect size vector.

$\epsilon\sim N(0,\sigma_{\epsilon}^{2}I)$ is the noise.

The solution of u for BLUP is given by (ref. [1]):

$$\hat{u}_{BLUP}=\sigma_{u}^{2}Z^{T}\left( Z\sigma_{u}^{2}Z^{T}+I\sigma_{\epsilon}^{2} \right)^{-1}\left( y-X \hat{\beta} \right)$$

Ridge regression is a simplified case of BLUP, with Z=X and $u$ **=** $\beta$:

$$y=X\beta+\epsilon$$

$$\hat{\beta}_{ridge}=\arg\min_{\beta} \left( \left\| y-X\beta\right\|^{2}+\lambda\left\| \beta\right\|^{2} \right)$$

The solution for $\beta_{ridge}$ can be analytically solved by:

$$\hat{\beta}_{ridge}=\left( X^{T}X+\lambda I \right)^{-1}X^{T}y$$

For BLUP, under the same simplified model, the solution of $\beta_{BLUP}$ is:

$$\hat{\beta}_{BLUP}=\left( X^{T}X+\frac{\sigma_{\epsilon}^{2}}{\sigma_{\beta}^{2}}I \right)^{-1}X^{T}y$$

Where $\sigma_{\beta}$ stands for the standard variance of the fixed effect size. It’s now easy to see BLUP and ridge regression are related under this simplified model, when

$$\lambda=\frac{\sigma_{\epsilon}^{2}}{\sigma_{\beta}^{2}}.$$

Since ZhangBLUP used this simplified model, it is equivalent to a ridge regression with a calculated $\lambda$ (ref. [2]).

#### PCA-preprocessed linear regression is mathematically related to ridge regression

A complete version of the below proof is proposed in ref. [3].

PCA performs a singular value decomposition on the coefficient matrix as:

$$X=UDV^{T}$$

The solution of a simple linear regression model $\hat{\beta}$ done on the PC-transformed coefficient matrix (A=UD) has below property:

$$A\hat{\beta}= UD\left( D^{T}U^{T}UD \right)^{-1}D^{T}U^{T}y=UU^{T}y$$

When the PCA incorporates PC selection, it becomes:

$$A\hat{\beta}= ULU^{T}y$$

Where L is a diagonal selection matrix with form

$$L=\left( \begin{matrix} \begin{matrix} 1 & & \\ & 1 & \\ & & \ddots\end{matrix} & & \\ & 1 & \\ & & \begin{matrix} 0 & & \\ & \ddots& \\ & & 0 \end{matrix} \end{matrix} \right),$$

where each 1 corresponds to a “selected” PC and 0 to a “ignored” PC.

For the solution of ridge regression:

$$X\hat{\beta}= X\left( X^{T}X+\lambda I \right)^{-1}X^{T}y=UD\left( D^{2}+\lambda I \right)^{-1}DU^{T}y$$

This form is close to PCA-preprocessed linear regression with

$$L=D\left( D^{2}+\lambda I \right)^{-1}D=\left( \begin{matrix} \frac{d_{1}^{2}}{d_{1}^{2}+\lambda} & & \\ & \frac{d_{p}^{2}}{d_{p}^{2}+\lambda} & \\ & & \begin{matrix} \ddots& \\ & \frac{d_{p}^{2}}{d_{p}^{2}+\lambda} \end{matrix} \end{matrix} \right),$$

where $d_{i}$ are the values along the diagonal line of the singular value matrix D in PCA. This observation implies that the ridge regression can be interpreted as a generalized, “smoothed” form of PCA-preprocessed linear regression, where principal component selection is not restricted to binary (0 or 1) choices. Taken together, these insights strongly suggest that the RD reduction in classic linear models is largely attributed by L2 regularization, implying that ridge-regression-like models may emerge as a promising technique for improving RD performances.

### Appendix

#### Public datasets

To ensure reproducibility and external validation, we included 11 publicly available datasets. GSE55763 (N = 2711, Illumina HumanMethylation 450K) [9] was downloaded from the Gene Expression Omnibus (GEO) database for model and PCA training for its large size, and was also used to assess age prediction RD using the 72 replicated samples. GSE83944 (N = 48, Illumina HumanMethylation 450K) [10], GSE247193 (N = 23 including one subject without replicate, Illumina Infinium EPIC Human Methylation Beadchip) [11], GSE247195 (N = 23 including one subject without replicate, Illumina Infinium EPIC Human Methylation Beadchip) [11] and GSE247197 (N = 46, Illumina HumanMethylation 450K) [11] were downloaded from the GEO database to assess age prediction RD as they all contain technical replicates. HTS-based datasets GSE144691 (N = 29) [12] and GSE86832 (N = 4) [13] were downloaded from the GEO database for assessment of unreliable CpGs, because they lacked replicates. Two additional datasets from the GEO database were downloaded for assessment of age prediction MAE and RD, namely GSE232346 (N = 24, TimeSeq) [14] and GSE245628 (N = 24, Illumina Infinium EPIC Human Methylation Beadchip) [14]. Finally, we also downloaded the Buccal cohort (Shokhirev & Johnson, 2025, N = 100, Twist and Illumina Infinium EPIC Human Methylation Beadchip v2) [15] and GSE164600 (N = 24, cfDNA WGBS) [16] datasets for various validation steps. The ethnicity and sex information can be found in the source dataset but was not considered in this study.

### Supplementary Tables

Table S1: Summary of epigenetic clocks for human age prediction investigated in this study. (Data provided in a separate file)

Table S2: Metadata of the 6 datasets generated in the study and SRRSH-141 cohort. (Data provided in a separate file)

Table S3: Percentage of clock CpGs with unreliable beta-values across HTS-based datasets. (Data provided in a separate file)

Table S4: Features and model coefficients of ClinicalAge. (Data provided in a separate file)

### Supplementary Figures

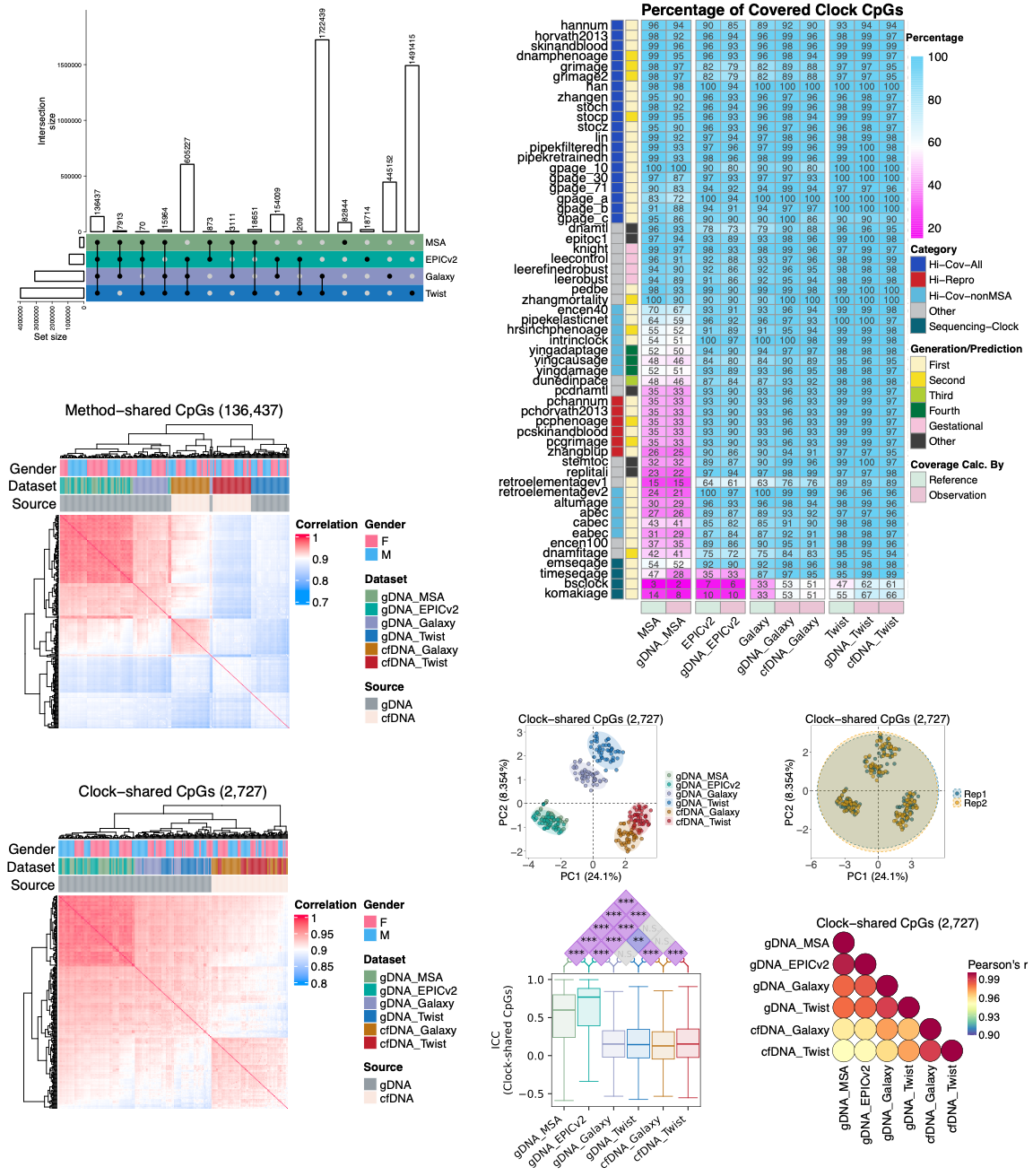

a

b

c

d

e

f

g

h

**Fig. S1| Precision and consistency estimated by clock CpGs were reduced in sequencing-based datasets. a**, An UpSet plot visualizing the intersections of CpG sites covered by the 4 different methods (MSA, EPICv2, Galaxy, and Twist). **b,** A heatmap displaying the coverage percentage of CpGs for 57 epigenetic clocks. Coverage is shown based on both the theoretical design of the 4 methods (manufacturer's specifications) and the empirical observations in 6 SRRSH-24 datasets (gDNA_MSA, gDNA_EPICv2, gDNA_Galaxy, gDNA_Twist, cfDNA_Galaxy, and cfDNA_Twist). The clocks are categorized by generation, predicted output, theoretical coverage, and observed reproducibility, as detailed in Fig. 1g. **c-d,** Principal Component Analysis (PCA) of all samples based on the “Clock-shared CpGs”, defined as the union of all CpGs from the “Hi-Cov-All” clocks. Samples are colored by dataset in (**c**) and by technical replicate in (**d**). **e-f,** Heatmaps showing pairwise Pearson correlation between samples. Correlations were calculated using beta-values from (**e**) “Method-shared CpGs” (N = 136,437), which are common to all four platforms, and (**f**) “Clock-shared CpGs”. Samples are annotated by gender, dataset, and DNA source. Hierarchical clustering was performed using Euclidean distance and complete linkage. **g,** Comparison of ICCs for Clock-shared CpGs across the 6 datasets. P-values were calculated by a two-tailed block permutation test (10,000 permutations) and adjusted for multiple comparisons using the Benjamini-Hochberg method. ***, p < 0.001; N.S., not significant. The center line of each boxplot marks the median, the box body marks the 1^st^ and 3^rd^ quartiles, and the caps mark the min and max values. **h,** A pairwise Pearson correlation matrix comparing the six datasets. Correlations were calculated using the mean beta-values of Clock-shared CpGs, where the mean was derived by averaging across all samples within each dataset.

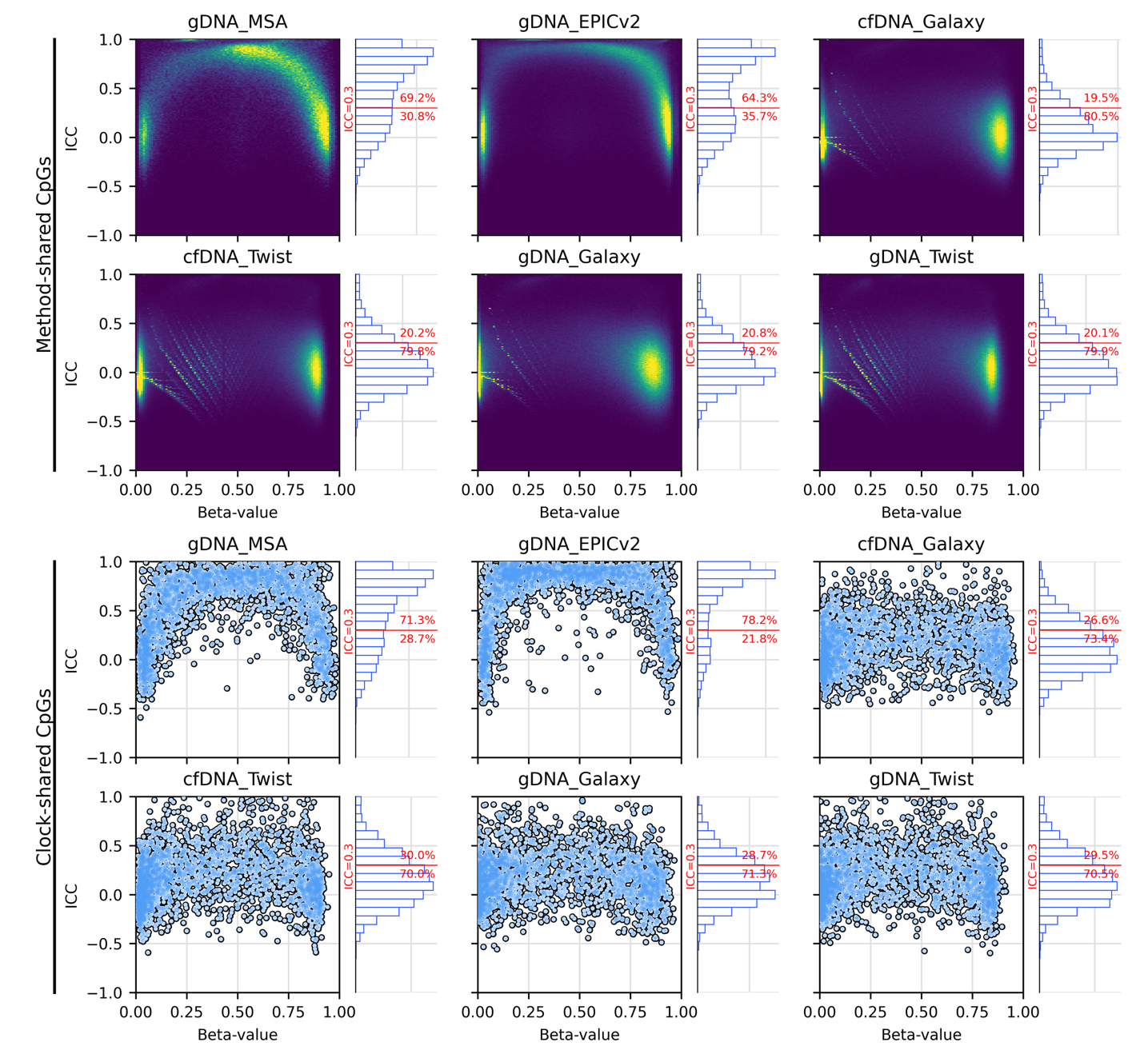

b

a

**Fig. S2| CpGs with low Interclass Correlation Coefficients (ICCs) were enriched around the 0 and 1 beta-values observed in all six SRRSH-24 datasets.** **a,** Summarized from 136,437 Method-shared CpGs; **b,** Summarized from 2,727 Clock-shared CpGs. ICC values were calculated by comparing the beta-values between replicates (intraclass) and across subjects (interclass). The beta-values on each plot represent the mean beta-values of each CpG locus.

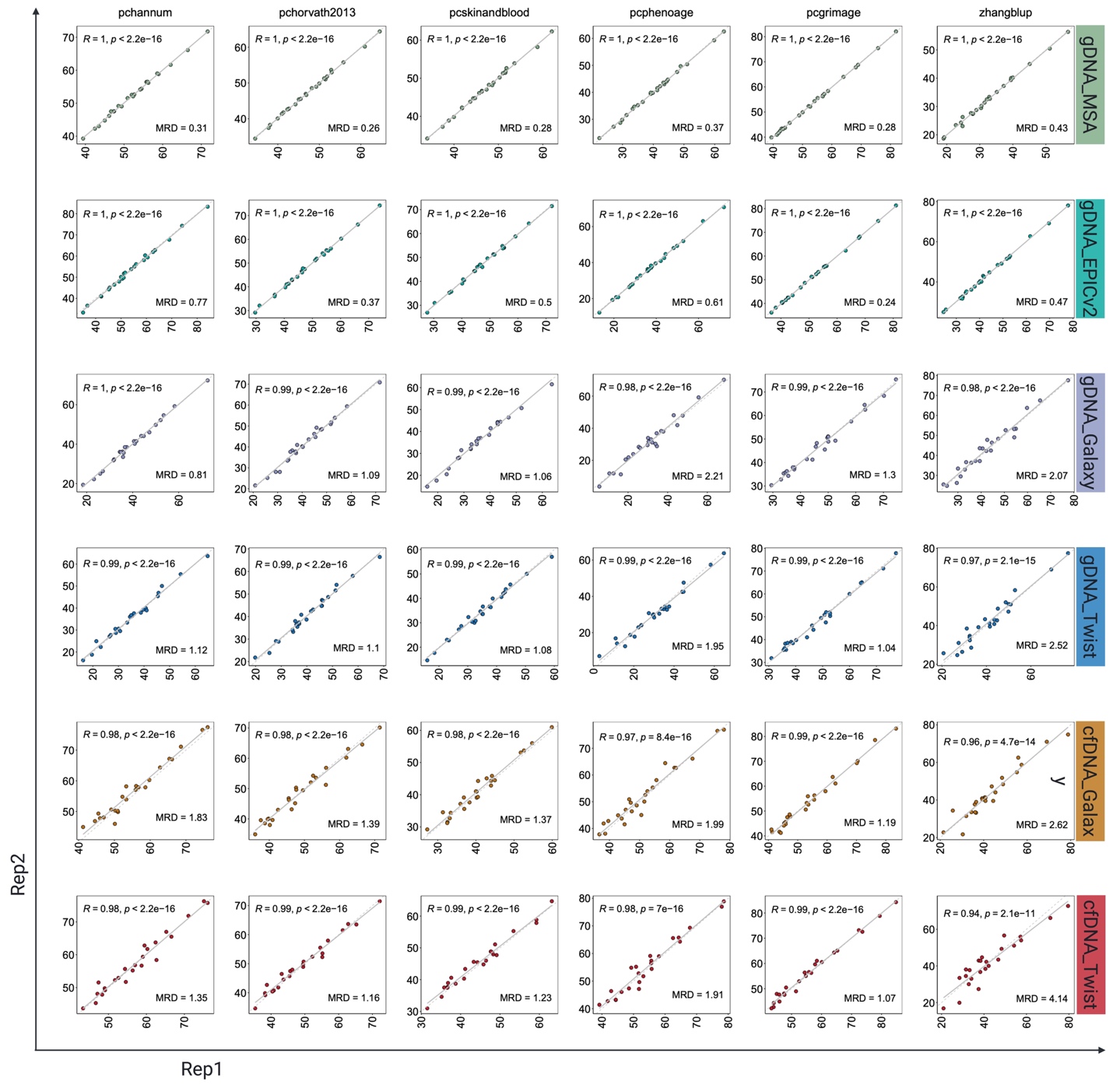

**Fig. S3| Hi-Repro clocks exhibited high reproducibility in all 6 SRRSH-24 datasets.** This figure consists of 6 panels, each dedicated to one of the 6 clocks. Within each panel, a scatter plot assesses the reproducibility of predicted ages by comparing technical replicates (Replicate 1 vs. Replicate 2). Each point represents an individual sample and is colored according to its source dataset. The plots include a line of identity (y=x, gray dashed line) and a linear regression fit (gray solid line). For each comparison, the Pearson’s R, p-value (N = 24), and the Mean Absolute Replicate Difference (MRD) are displayed directly on the panel.

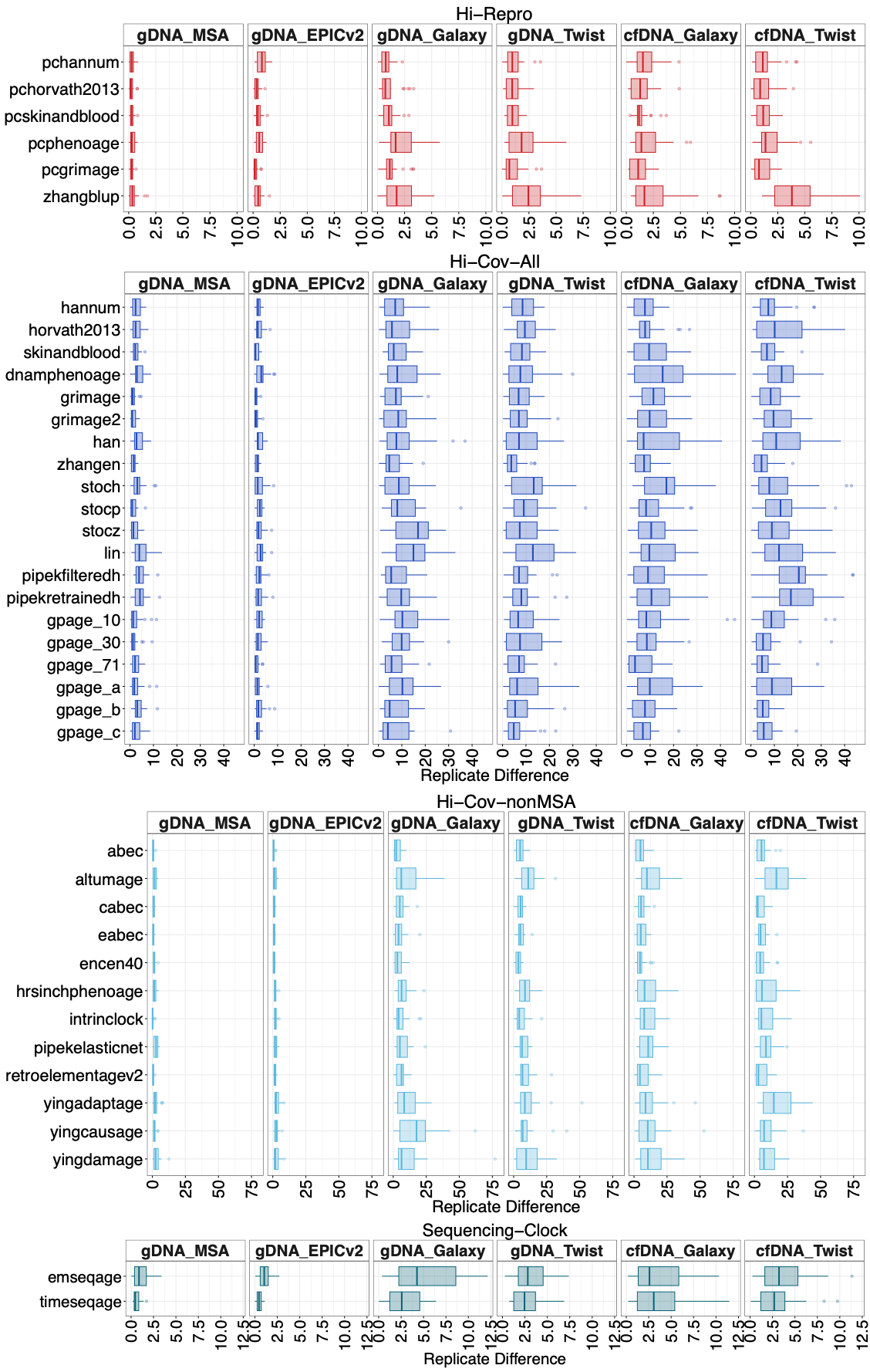

d

c

b

a

**Fig. S4| Reduced reproducibility of epigenetic clocks in HTS-based datasets.** Boxplots comparing the distribution of inter-replicate differences in predicted age across the 6 SRRSH-24 datasets (N=24 samples per dataset). The difference is calculated as the absolute difference between two technical replicates. The analysis is presented for three distinct categories of epigenetic clocks: **a,** Hi-Repro clocks; **b,** Hi-Cov-All clocks; **c,** Hi-Cov-nonMSA clocks; and **d,** Sequencing clocks. A wider distribution indicates lower reproducibility. The center line of each boxplot marks the median, the box body marks the 1^st^ and 3^rd^ quartiles, and the whisker stretches to the min and max values.

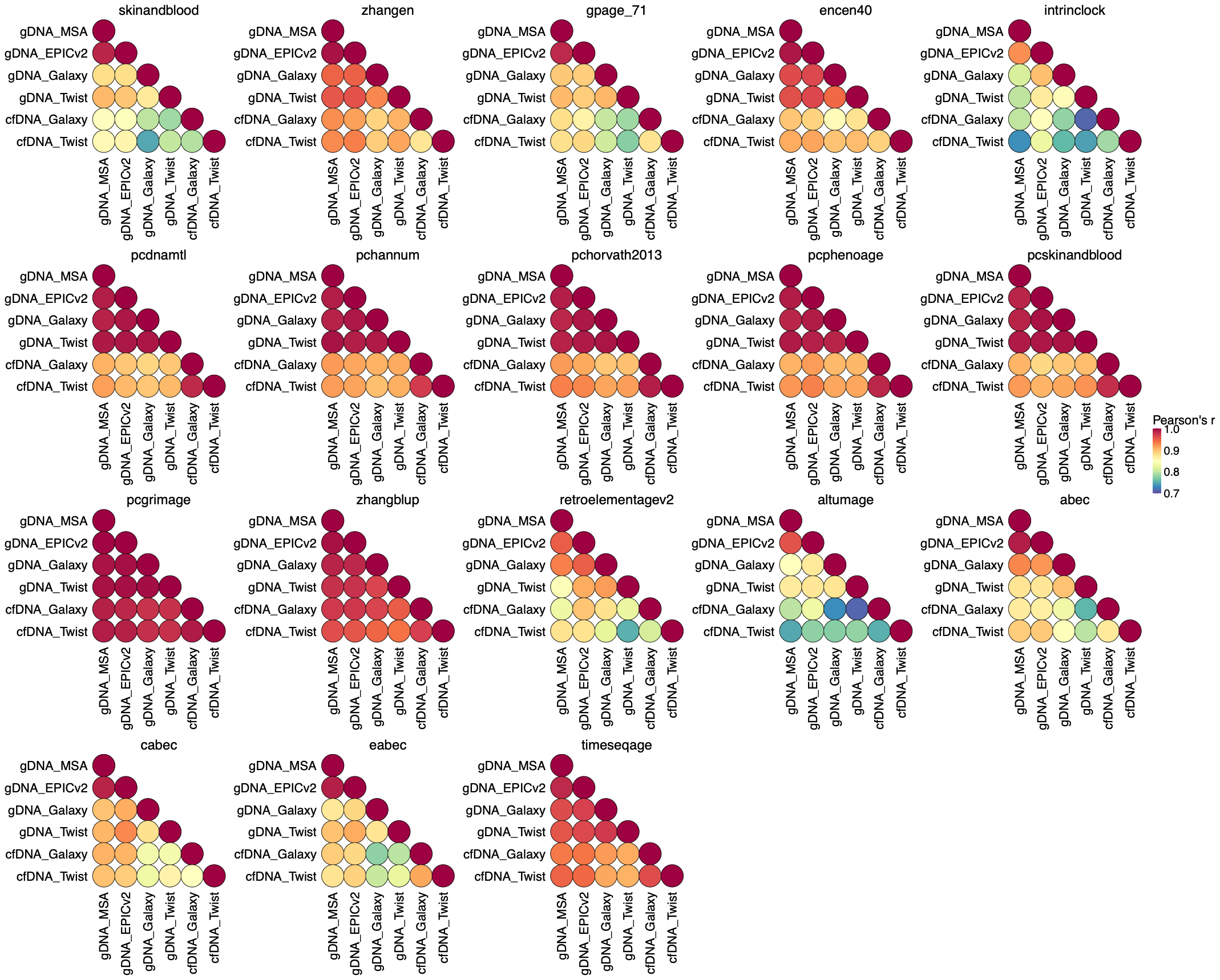

**Fig. S5| High cross-dataset consistency for 18 selected epigenetic clocks.** The figure displays a heatmap of pairwise Pearson’s R, assessing the consistency of predicted ages for each clock across all datasets. To perform this analysis, the predicted ages from the two technical replicates for each sample were first averaged. The correlations were then calculated between each pair of datasets using these mean age values (N = 24). The 18 clocks shown are those that exhibited R ≥ 0.7 in all an average cross-dataset comparisons.

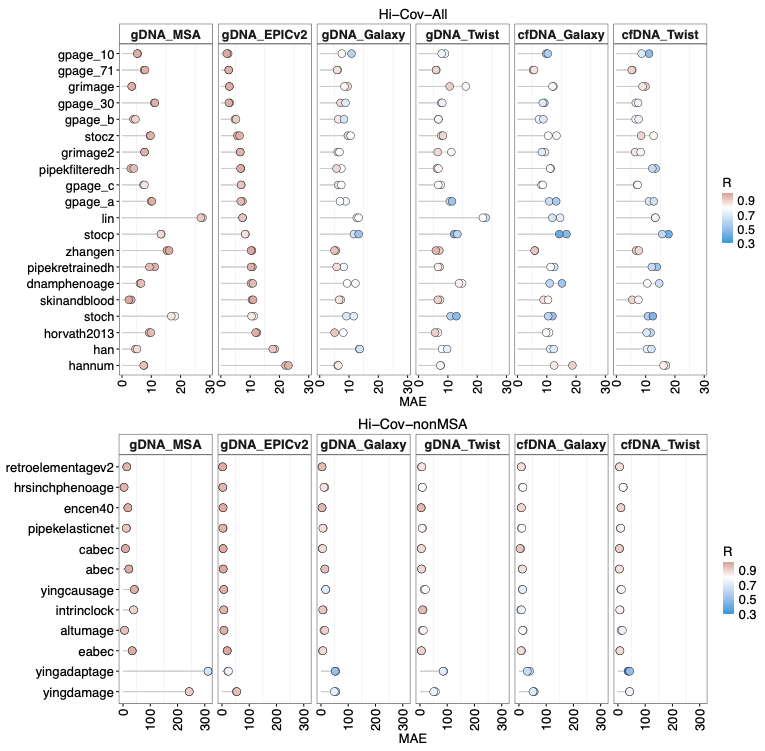

b

a

**Fig. S6| Reduced epigenetic clock performance in HTS-based datasets, tested with Hi-Cov-All and Hi-Cov-nonMSA clocks. a,** A lollipop plot illustrating the performance and variability of 20 Hi-Cov-All clocks. **b,** A lollipop plot illustrating the performance and variability of 12 Hi-Cov-nonMSA clocks. For each clock, the two points represent the Mean Absolute Error (MAE) of the two technical replicates against chronological age. The color of the points corresponds to Pearson’s R between predicted and chronological age. The clocks are sorted in ascending order based on the average MAE observed in the gDNA_EPICv2 dataset.

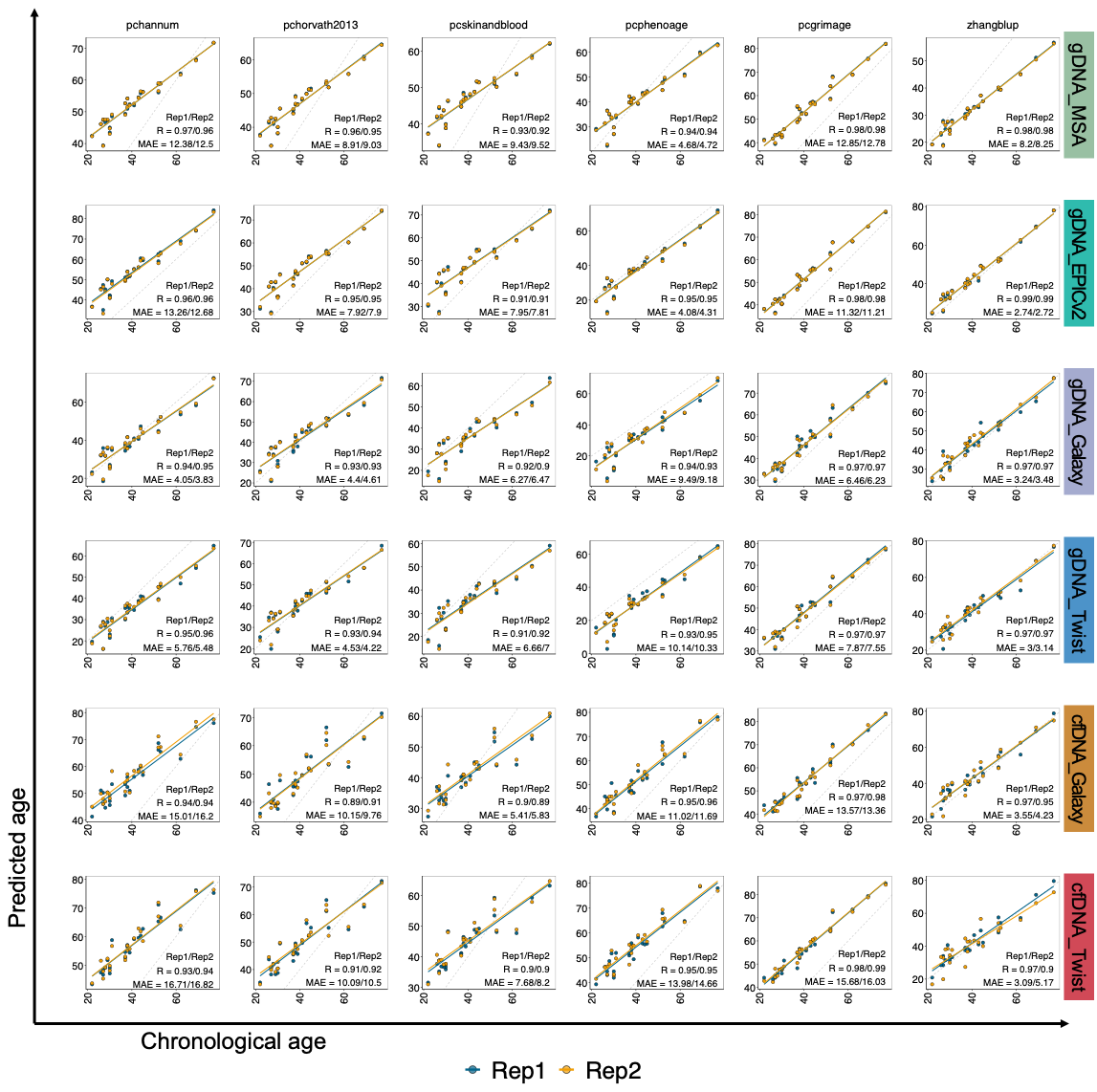

**Fig. S7| Age prediction accuracy analysis of Hi-Repro clocks by replicate groups.** For each clock and replicate, the predicted epigenetic age (y-axis) is plotted against the chronological age (x-axis). The figure includes the line of identity (y=x, gray dashed line) and a linear regression fit (colored solid line). The corresponding Pearson’s R and MAE values are displayed on each panel.

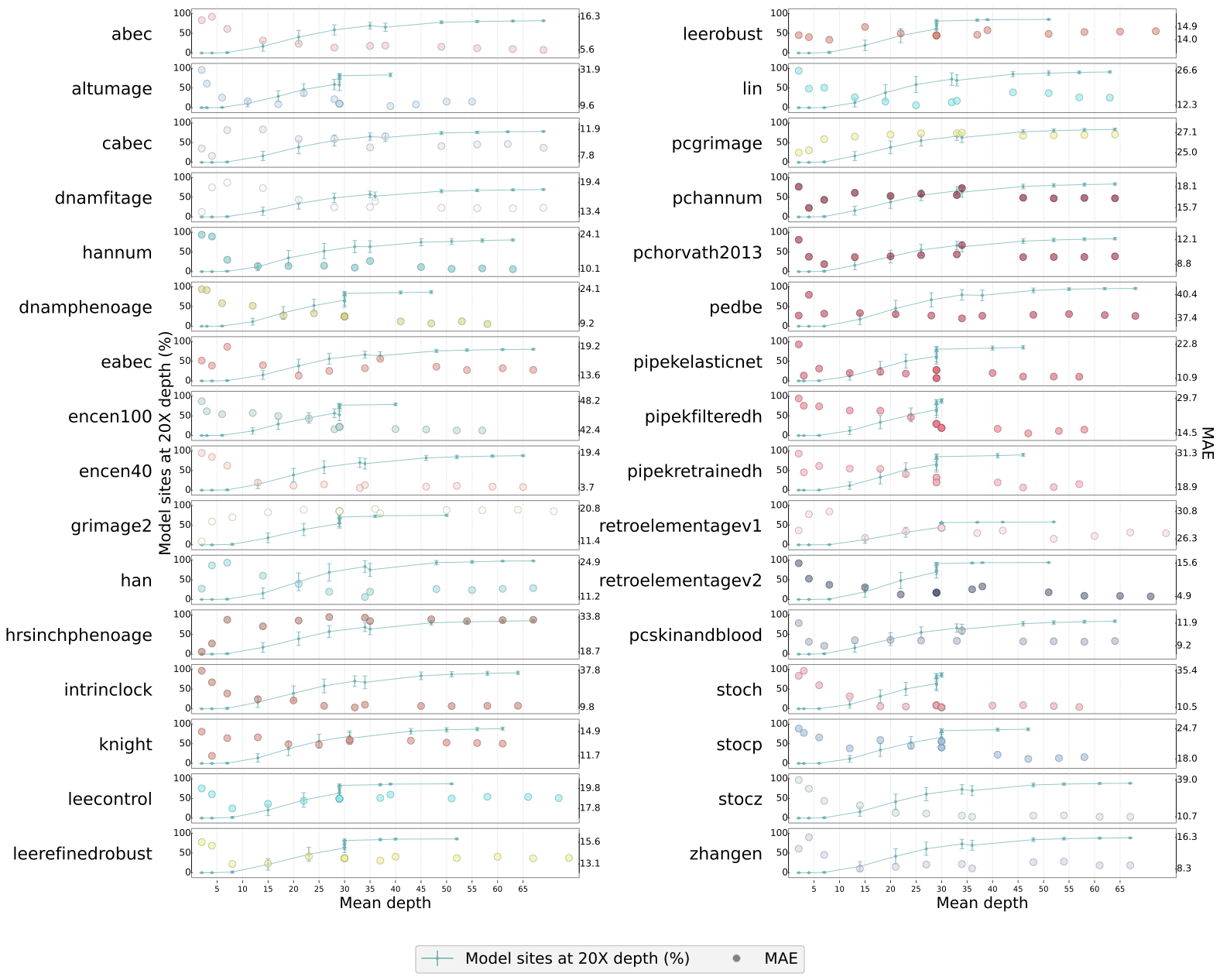

**Fig. S8| Heterogeneous sequencing depth requirements for epigenetic clock performance.** This figure illustrates the performance of various epigenetic clocks as a function of sequencing depth, based on a downsampling analysis. Each panel contains a line plot showing the Mean Absolute Error (MAE) for a specific clock, calculated from datasets that were computationally downsampled to different mean coverage levels. The analysis reveals that clocks have distinct requirements for sequencing depth to achieve stable performance. Two general patterns were observed: (1) One group of clocks, including hannum, zhangen, stoch, stocz, lin, pedbe, and encen40, achieves a stable MAE at a mean sequencing depth of approximately 20×; and (2) A second, more data-demanding group of clocks, including han, pipekfilteredh, pipekretrainedh, pipekelasticnet, intrinclock, altumage, abec, and encen100, requires a minimum depth of 30× or greater for their performance to converge.

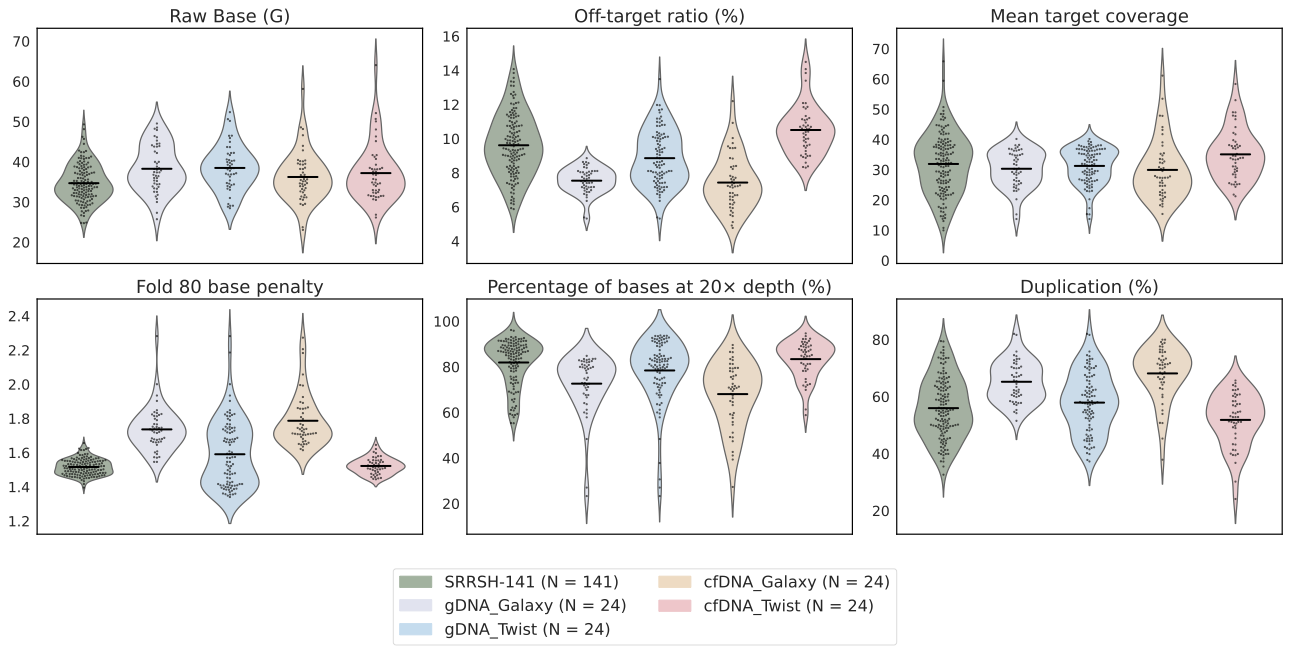

**Fig. S9| Primary Quality Control (QC) metrics for the study cohorts (SRRSH-24 and SRRSH-141).** This figure displays a series of violin plots comparing key QC metrics across the four HTS-based datasets from the SRRSH-24 cohort and the SRRSH-141 validation cohort. The SRRSH-24 datasets include technical replicates, whereas the SRRSH-141 cohort does not. To ensure a fair comparison across all platforms and cohorts, all metrics shown (with the exception of Raw Bases) were calculated after downsampling the raw sequencing data for each sample to a uniform depth of 30 Gb. The horizontal black line within each violin plot indicates the mean value for that dataset.

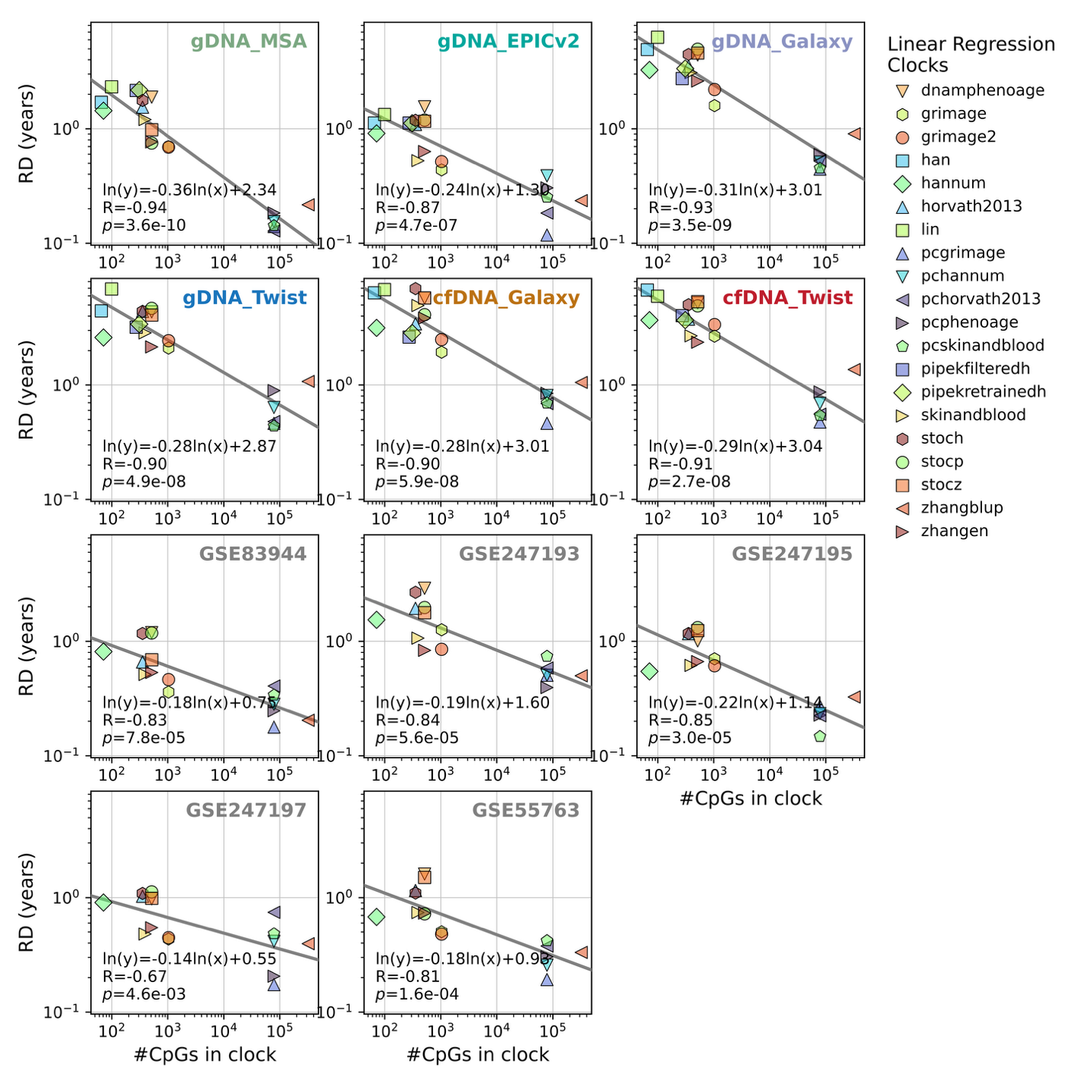

**Fig. S10| A consistent negative correlation exists between the number of CpGs in a clock and its prediction variability.** This figure presents a series of eleven scatter plots, one for each of the datasets analyzed, to test the hypothesis that clocks with more CpGs are more reproducible. In each plot, the x-axis represents the number of CpG sites composing a clock, and the y-axis shows the mean Replicate Difference (RD) in predicted age between technical replicates. Each data point corresponds to one of 20 HCRL clocks. HCRL clocks denote 20 linear-architecture clocks classified as High-Cov-All or High-Repro. A negative correlation was consistently observed across all datasets, confirming that clocks built with a larger number of CpGs tend to have lower variability and higher reproducibility. The analysis was independently performed on six datasets from this study (gDNA_MSA, gDNA_EPICv2, gDNA_Galaxy, gDNA_Twist, cfDNA_Galaxy, and cfDNA_Twist) and five publicly available datasets (GSE83944, GSE247193, GSE247195, GSE247197, and GSE55763). All observed correlations were statistically significant after correcting for multiple comparisons using the Benjamini-Hochberg procedure (all adjusted p-values < 4.7 x 10^-3^).

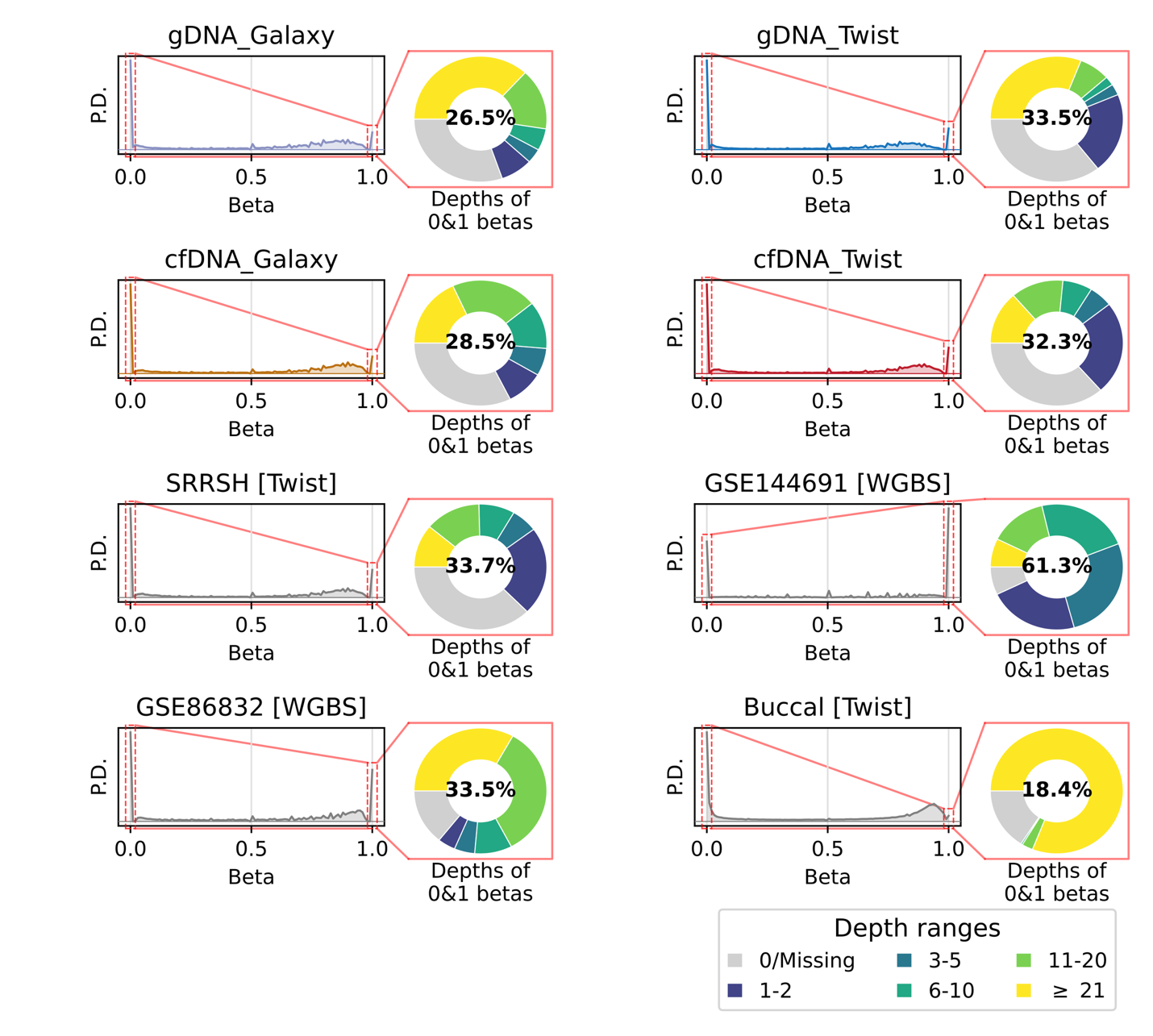

**Fig. S11| High prevalence of unreliable beta-values in HTS-based methylation datasets.** HTS-based methylation profiling technologies are susceptible to generating unreliable beta-values (exactly 0 or 1), which can account for up to 61.3% of all CpG sites in a given dataset. This figure presents a series of circular charts, one for each of the eight datasets analyzed (4 from this study and 4 publicly available). The percentage shown in the center of each circle indicates the proportion of total CpGs in that dataset with unreliable beta-values. The Buccal cohort (Shokhirev and Johnson) is shown for reference but was pre-processed with a filter that removed all data with a sequencing depth of 10 or less, resulting in the absence of low-depth observations for this sample.

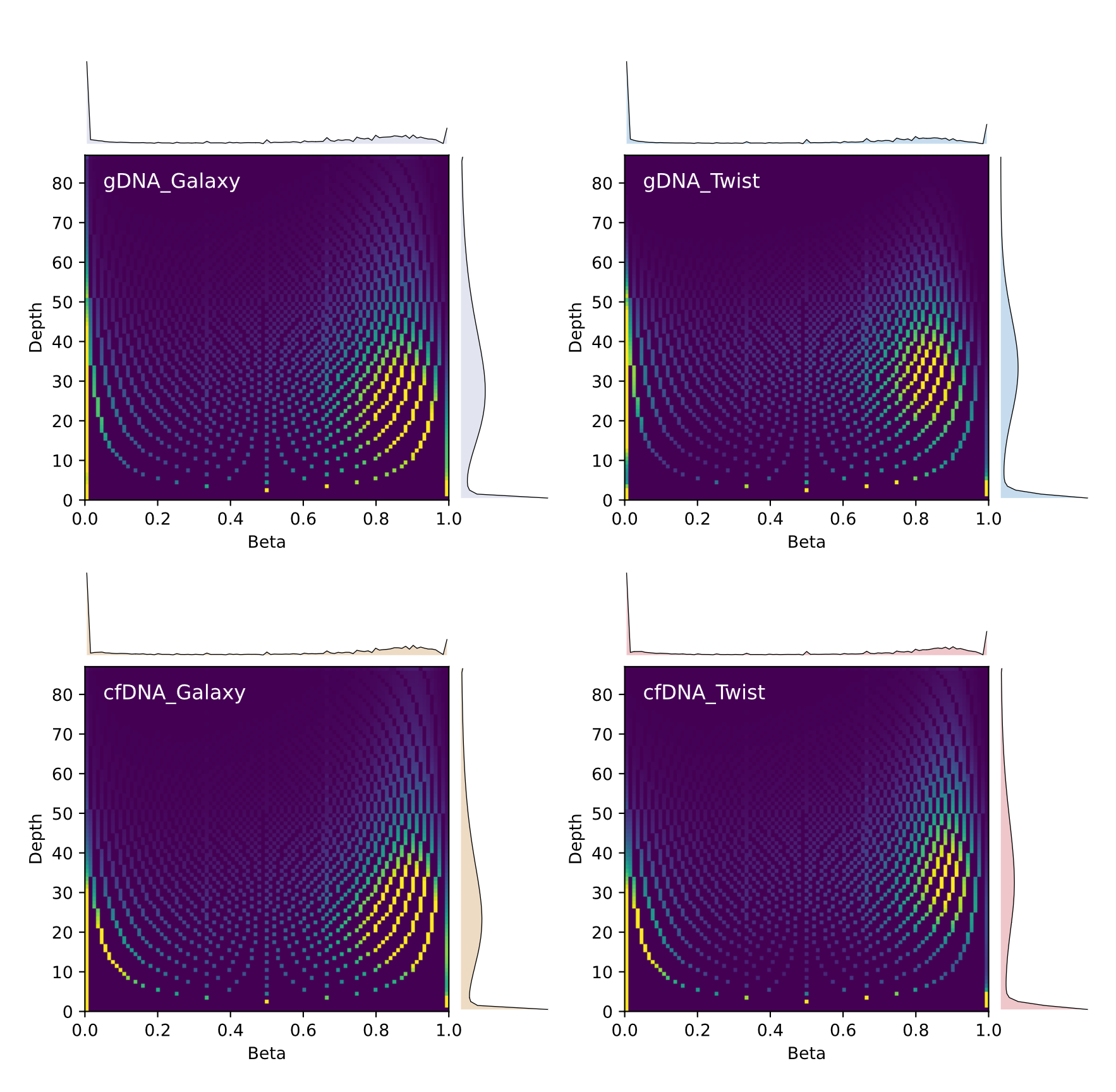

**Fig. S12| Joint and marginal distributions of beta-values and sequencing depth in HTS datasets.** This figure illustrates the relationship between beta-values and sequencing depth across four HTS-based datasets using joint and marginal distribution plots. For each dataset, the central heatmap shows the joint distribution as a 2D density plot, while histograms on the axes display the marginal distributions for depth and beta-values, respectively. A prominent high-density spike is observed at depth=0 and beta=0 in all datasets. This is an artifact resulting from the common practice of encoding missing data (uncovered CpG sites) with these values. Notably, the plots reveal the inherently discrete nature of sequencing-derived beta-values. At a given sequencing depth d, a beta-value is calculated from integer read counts and can thus only take on d + 1 possible fractional values. For example, at a sequencing depth of 2×, the only possible beta-values are 0, 0.5, and 1.0.

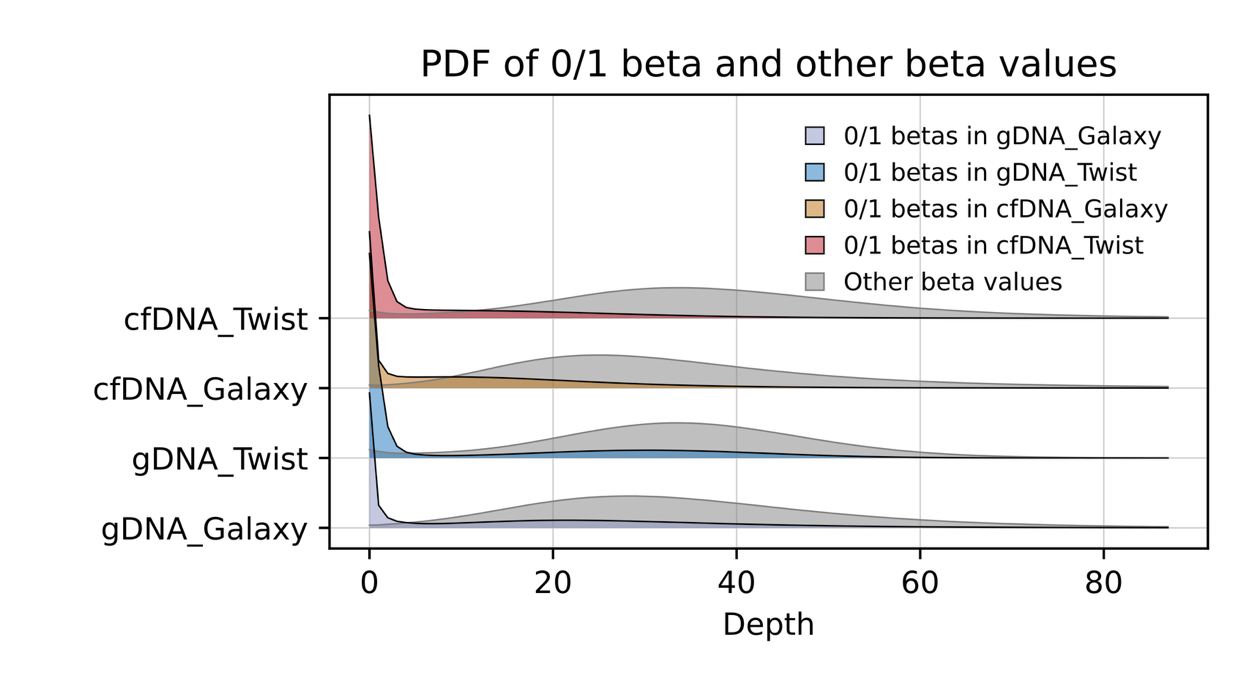

**Fig. S13| Unreliable and normal CpGs exhibit distinct sequencing depth distributions.** This figure compares the distribution of sequencing depth for two categories of CpGs: unreliable CpGs (defined as those with beta-values of exactly 0 or 1) and normal CpGs (with beta-values 0 < beta < 1). The overlaid density plots consistently demonstrate that unreliable CpGs are predominantly observed at very low sequencing depths. In contrast, normal CpGs show a much broader distribution shifted towards higher coverage. This finding strongly suggests that the majority of unreliable 0/1 beta-values are technical artifacts arising from insufficient sequencing coverage, rather than representing a true biological state of complete methylation or unmethylation. PDF: probability density function.

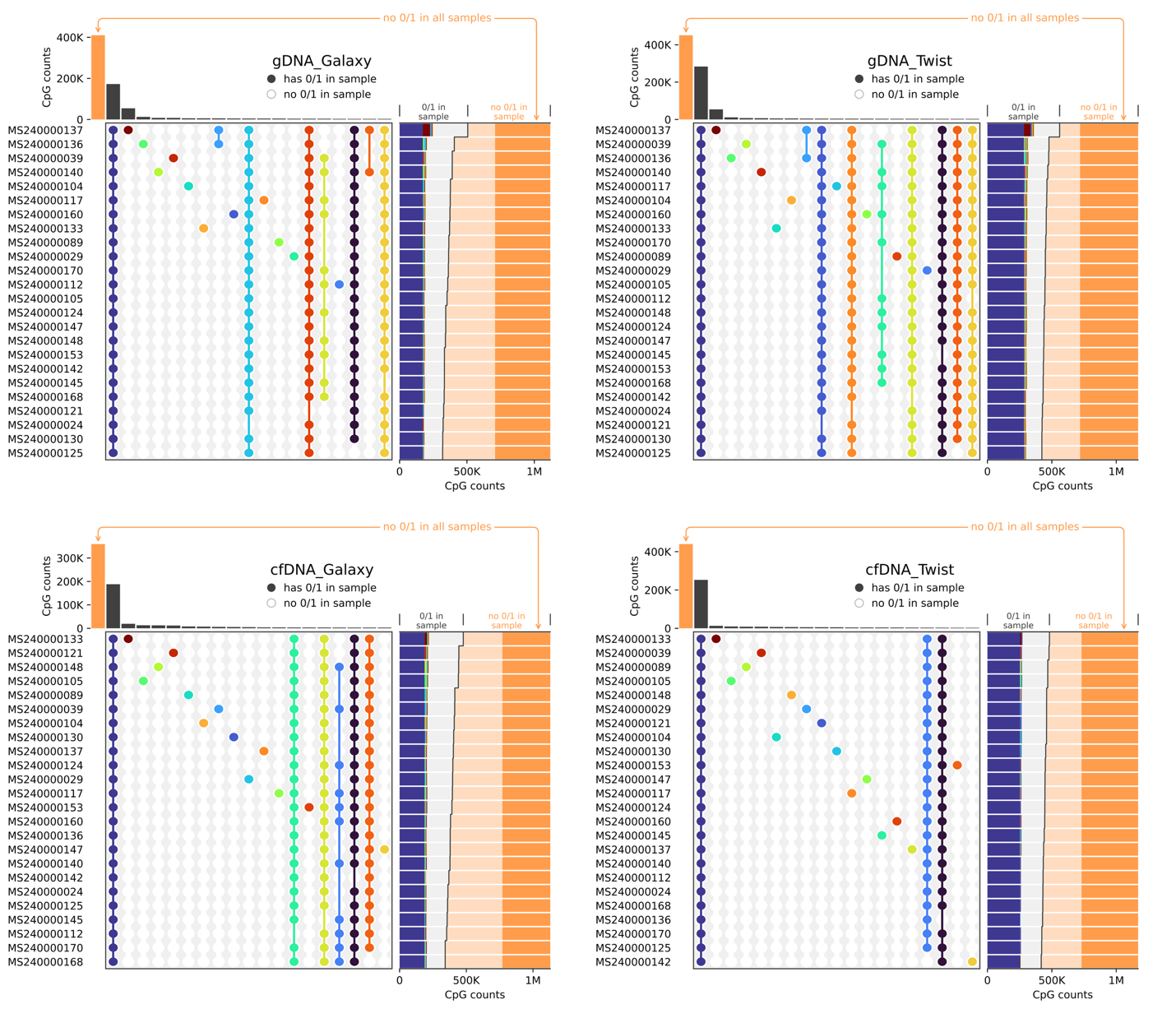

**Fig. S14| Distribution of unreliable CpGs across samples within 4 SRRSH-24 HTS datasets.** This figure presents a series of histograms, one for each HTS dataset generated in this study, illustrating the recurrence of unreliable CpGs across the 24 samples. The x-axis represents the number of samples in which a specific CpG site was flagged as unreliable, and the y-axis indicates the count of such CpGs. A large proportion of unreliable CpGs (ranging from approximately 30% to 70% depending on the dataset) are consistently unreliable across all 24 samples (highlighted in dark blue). This pattern suggests these are systematically uncovered sites, likely representing regions that failed to be captured or sequenced in that specific assay.

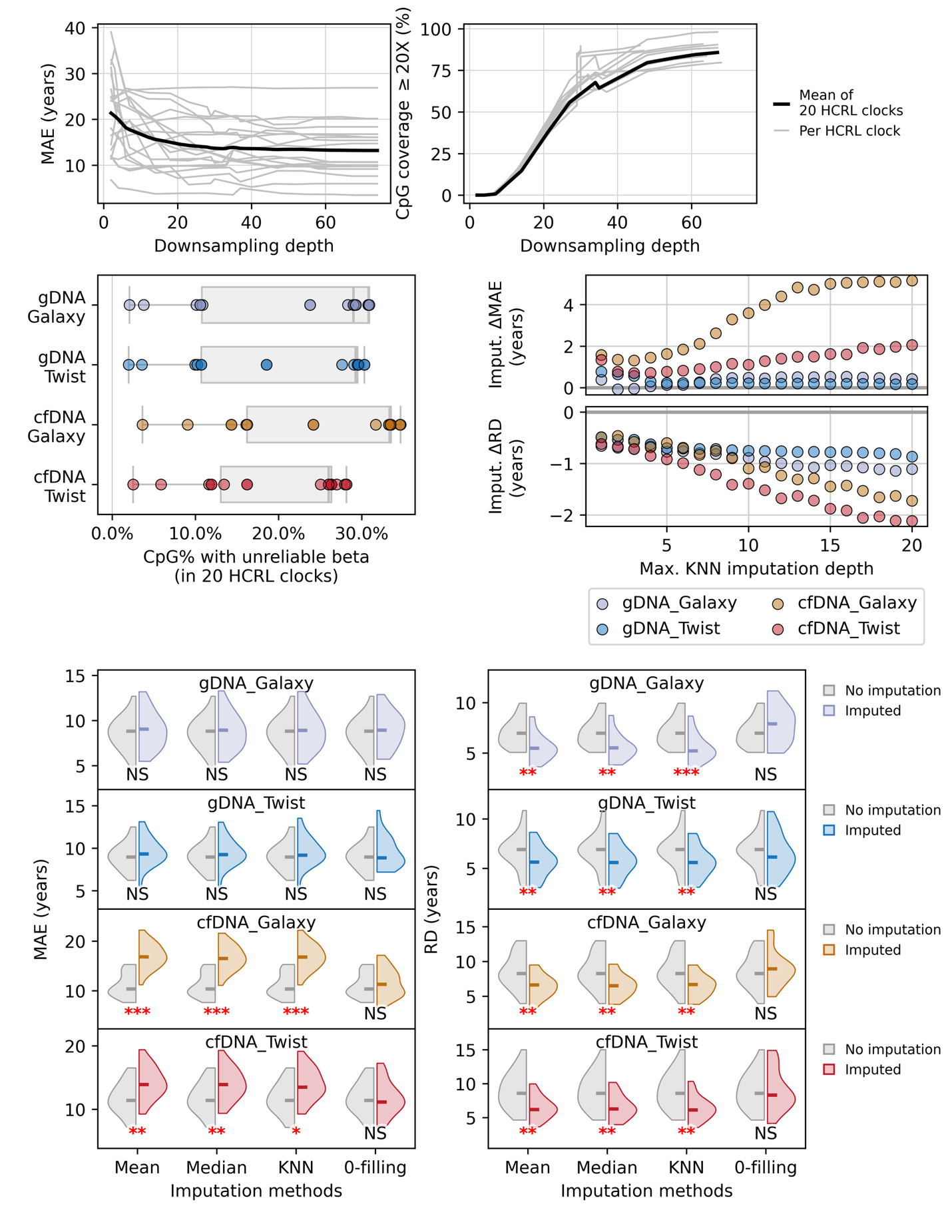

**Fig. S15| Benchmarking of age prediction improvements across four common imputation methods (mean, median, KNN and 0-filling).** The horizontal lines mark the medians of each respective distribution. No significant increase of MAE was observed in gDNA datasets. 0-filling yielded neither significant improvement nor deterioration in MAE or RD. P-values were obtained by two-sided permutation tests (10,000 permutations, custom script) comparing pre- and post-imputation predictions. Multiple testing results were adjusted using the Benjamini-Hochberg procedure. ***, p < 0.001; **, p < 0.01; *, p < 0.05; NS, not significant.

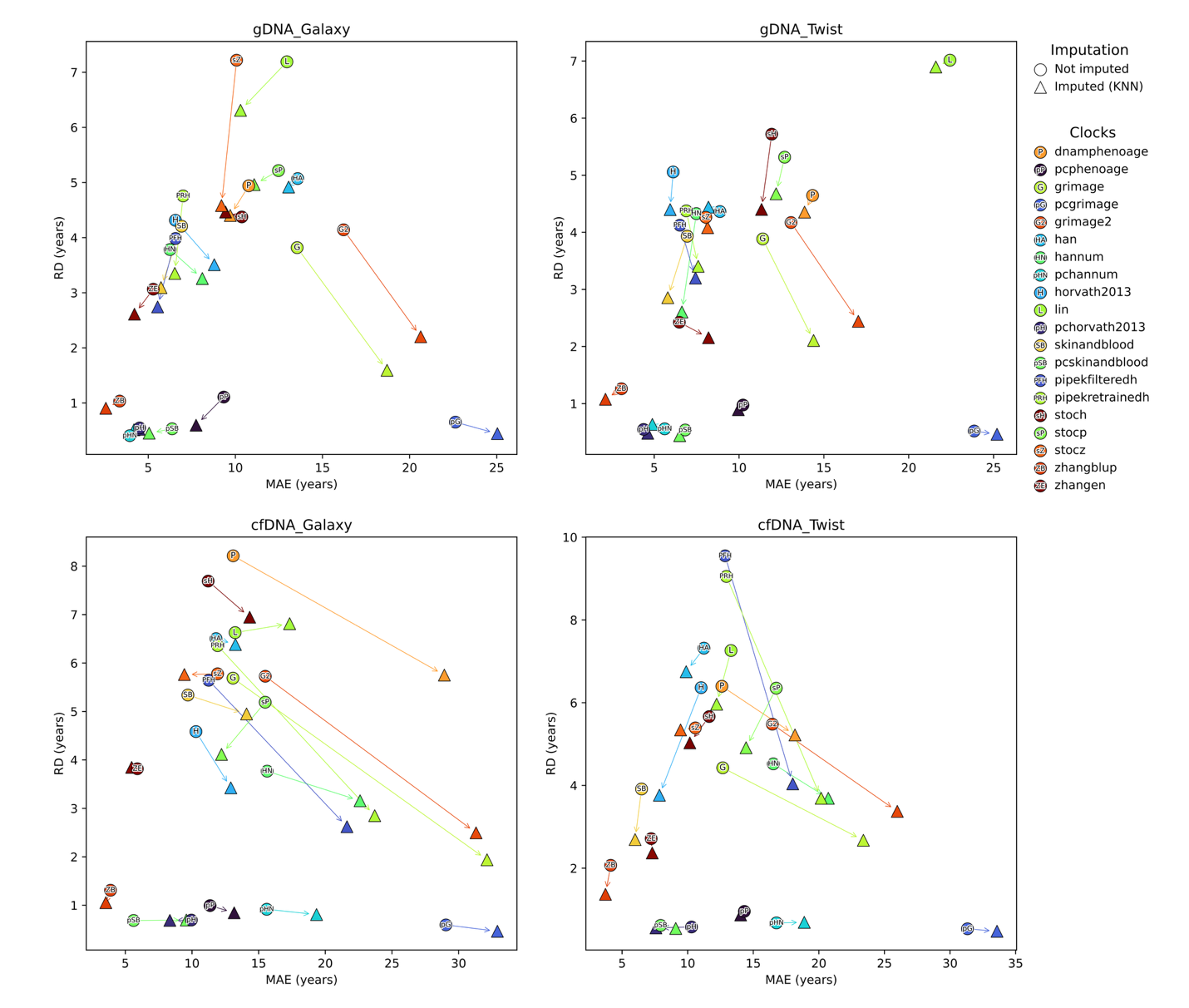

**Fig. S16| Effect of KNN imputation on the performance of 20 HCRL clocks.** This figure presents a series of scatter plots evaluating the impact of K-Nearest Neighbors (KNN) imputation on clock performance across the four HTS-based datasets. HCRL clocks denote 20 linear-architecture clocks classified as High-Cov-All or High-Repro. For each clock, the Mean Absolute Error (MAE) is plotted on the x-axis, and Replicate Difference (RD) is on the y-axis. The performance before and after the imputation is displayed as the movement in this MAE-RD plane. While the effects of imputation varied, only the zhangblup (ZB) and stocp (sP) clocks consistently demonstrated improved performance for both MAE and RD across all four datasets.

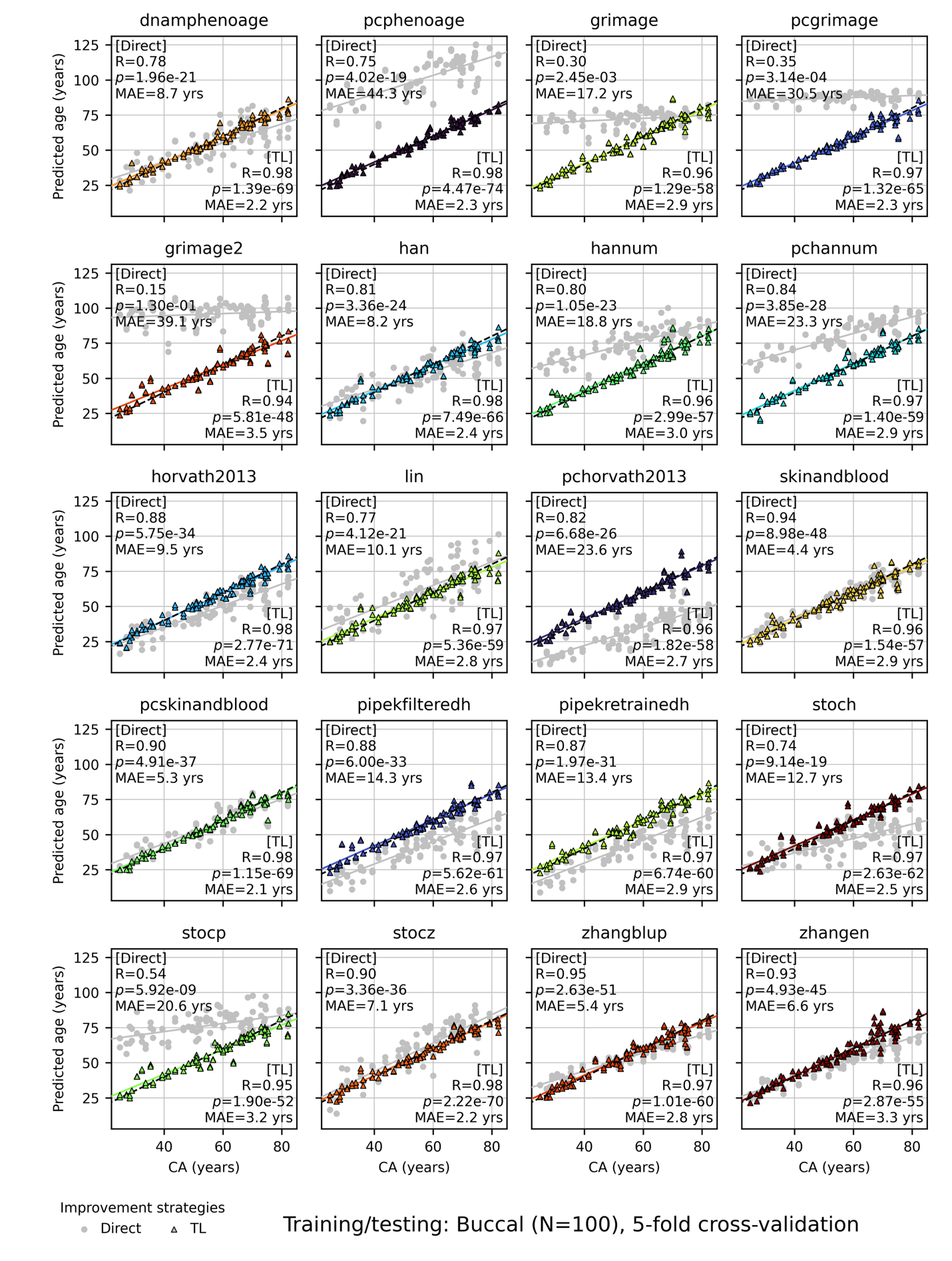

**Fig. S17| The prediction MAE and correlation (Pearson’s R) improvements achieved by distillation transfer learning (TL) in the Buccal cohort (N = 100).** The grey solid lines represent the linear regression lines between direct predictions (without TL) of each clock and chronological age (CA) calculated from clinical measurements. The colored solid lines represent the linear regression lines between TL predictions of each clock and CA. The black dashed lines mark the diagonal.

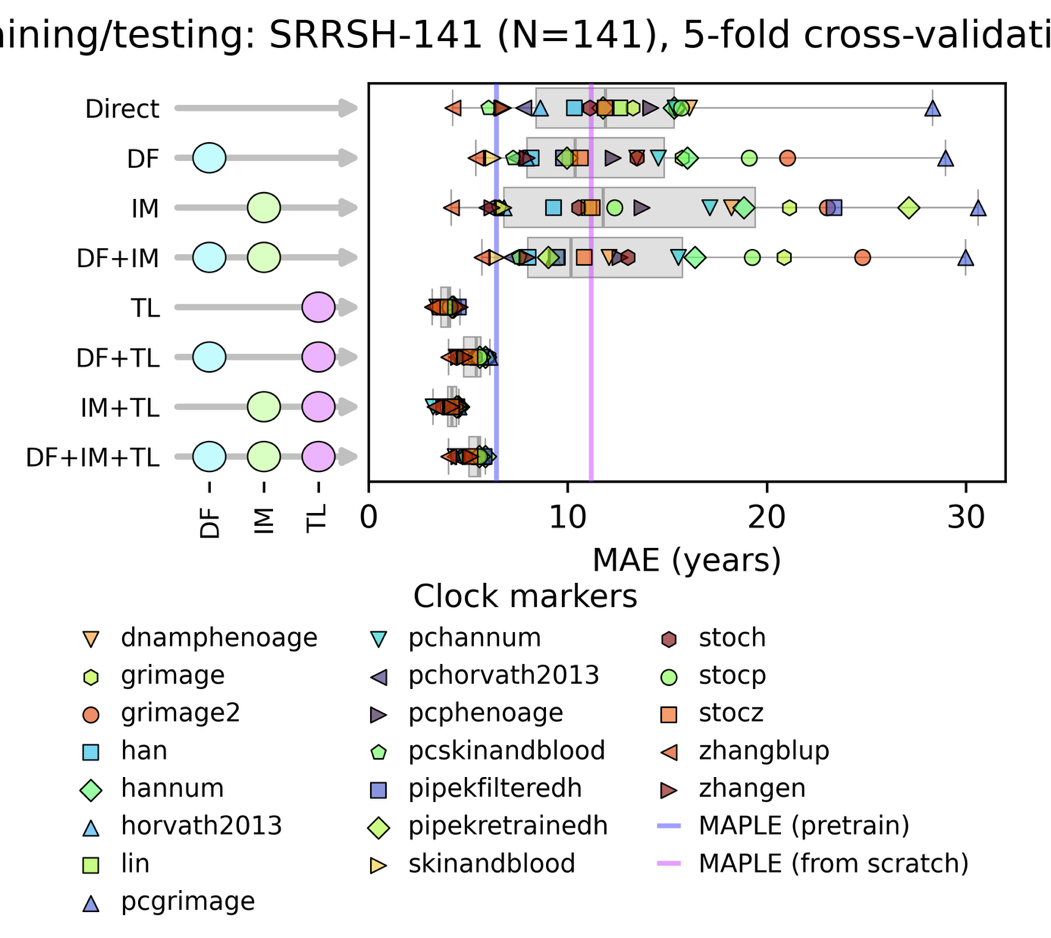

**Fig. S18| Evaluating strategies for improving epigenetic clock accuracy using the SRRSH-141 cfDNA cohort.** This figure compares the performance of 20 HCRL clocks on the SRRSH-141 cohort, evaluated under four distinct data processing strategies. HCRL clocks denote 20 linear-architecture clocks classified as High-Cov-All or High-Repro. The results are displayed as a series of boxplots, with each plot illustrating the distribution of Mean Absolute Error (MAE) from the 20 clocks for a given strategy. The four strategies are defined as follows: **Direct**, Baseline performance calculated without any optimization techniques; **DF** (Depth Filtering), Filtering out CpG sites with a sequencing depth below 20×; **IM** (Imputation), Imputation of unreliable (0 or 1) beta-values; and **TL** (Transfer Learning), Applying distillation model. The center line of each boxplot marks the median, the box body marks the 1^st^ and 3^rd^ quartiles, and the caps mark the minimum and maximum values.

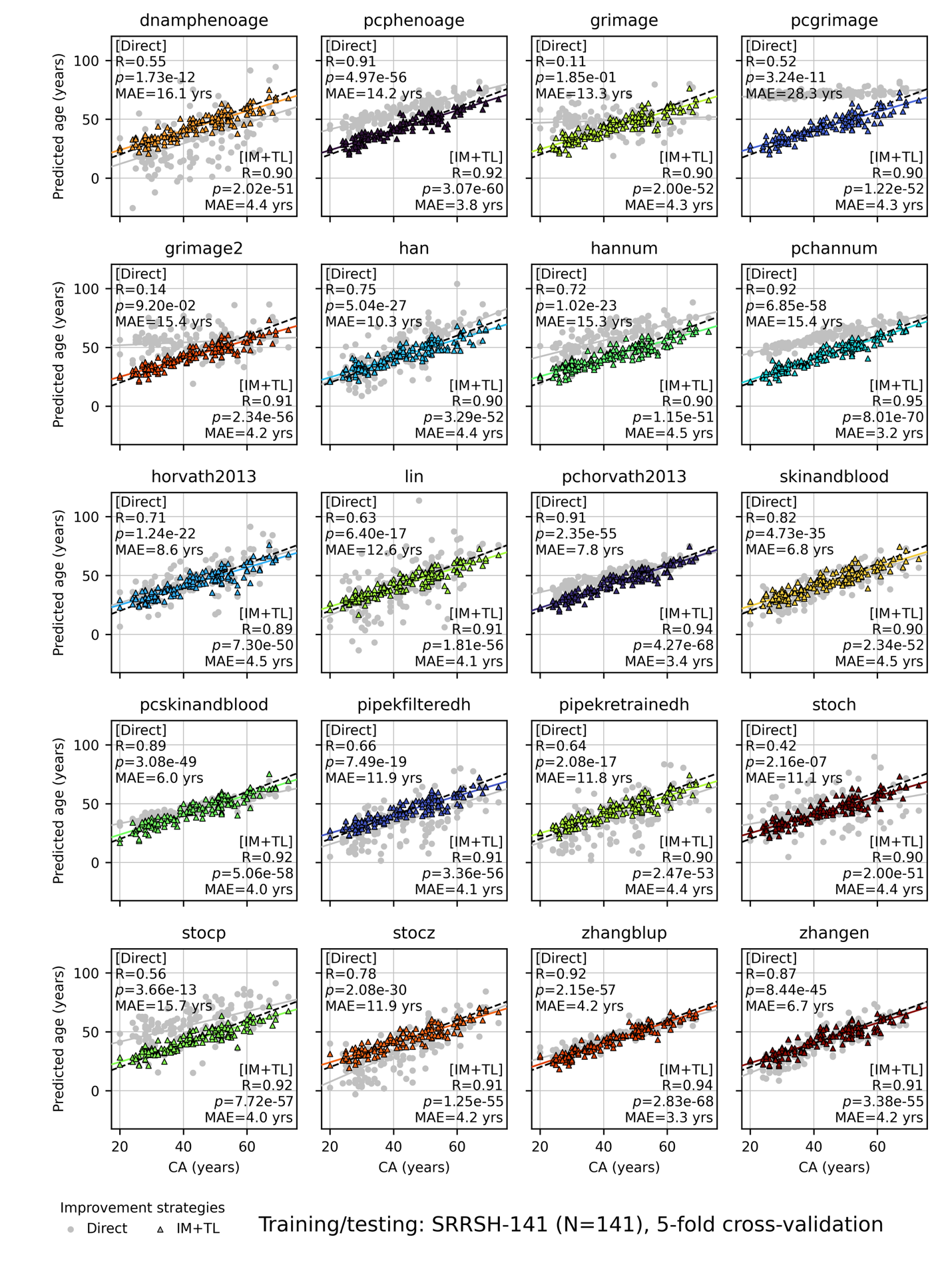

**Fig. S19| The prediction MAE and correlation (Pearson’s R) improvements achieved by combining unreliable beta imputation and distillation (IMputation+Transfer Learning, IM+TL) in the SRRSH-141 cohort (N = 141).** The grey solid lines represent the linear regression lines between direct predictions (without TL) of each clock and chronological age (CA) calculated from clinical measurements. The colored solid lines represent the linear regression lines between TL predictions of each clock and CA. The black dashed lines mark the diagonal.

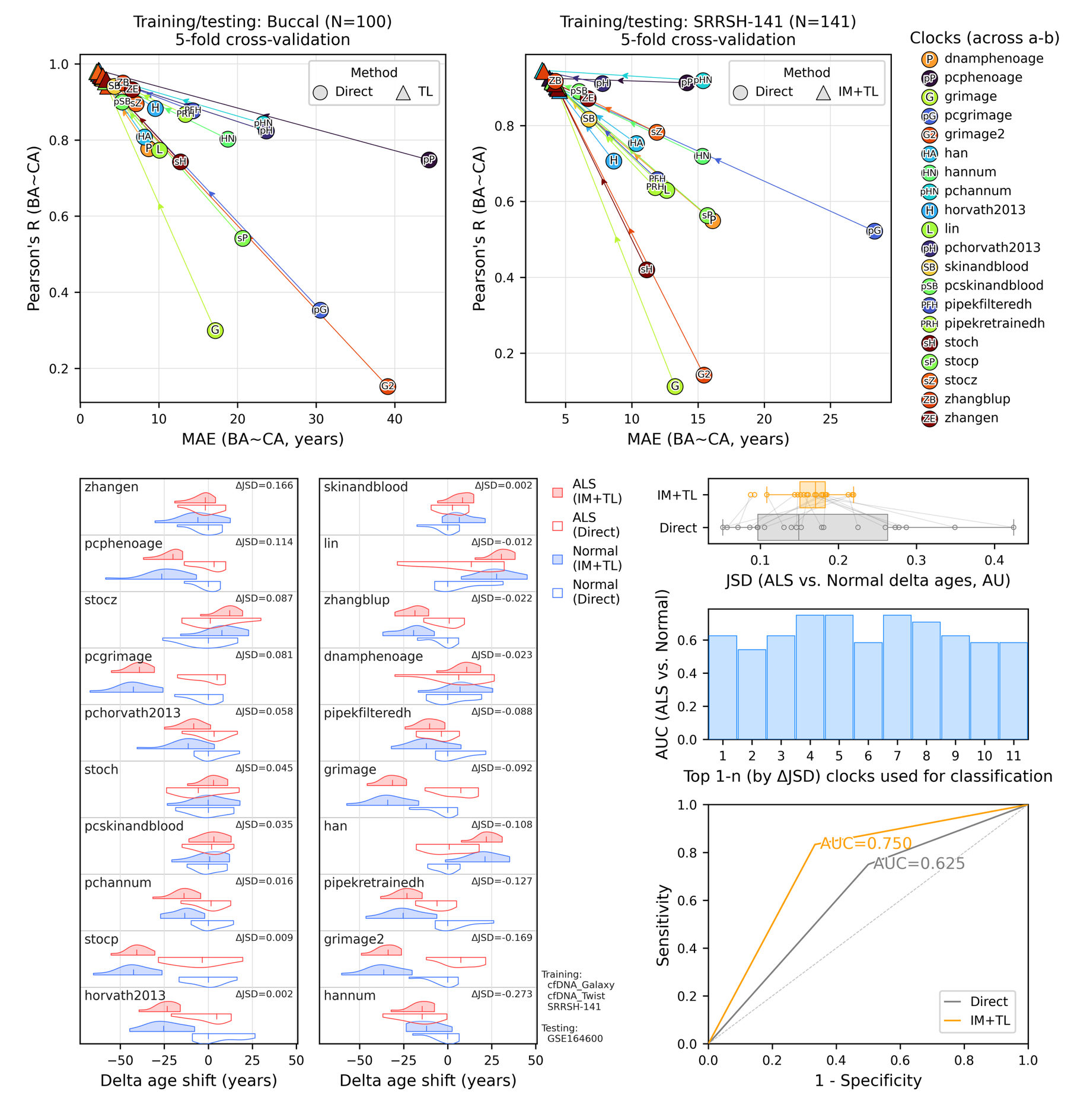

**Fig. S20|** Distributional change of delta-age predicted by 20 HCRL clocks before and after clock adaptation (IM+TL as the best strategy), performed on public dataset GSE164600. The center line in each violin plot marks the median.
